# Conservation of heat stress acclimation by the inositol polyphosphate multikinase, IPMK responsible for 4/6-InsP_7_ production in land plants

**DOI:** 10.1101/2023.11.17.567642

**Authors:** Ranjana Yadav, Guizhen Liu, Priyanshi Rana, Naga Jyothi Pullagurla, Danye Qiu, Henning J. Jessen, Debabrata Laha

## Abstract

Inositol pyrophosphates (PP-InsPs) are soluble cellular messengers that integrate environmental cues to induce adaptive responses in eukaryotes. In plants, the biological functions of various PP-InsP species are poorly understood, largely due to the absence of canonical enzymes present in other eukaryotes. The recent identification of a new PP-InsP isomer with yet unknown enantiomeric identity, 4/6-InsP_7_ in the eudicot *Arabidopsis thaliana*, further highlights the intricate PP-InsP signalling network employed by plants. The abundance of 4/6-InsP_7_ in land plants, the enzyme(s) responsible for its synthesis, and the physiological functions of this species are all currently unknown. In this study, we show that 4/6-InsP_7_ is the major PP-InsP species present across land plants. Our findings demonstrate that the *Arabidopsis* inositol polyphosphate multikinase (IPMK) homolog, AtIPK2α generates 4/6-InsP_7_ *in vitro*. Furthermore, the cellular level of 4/6-InsP_7_ is controlled by the two *Arabidopsis* IPMK isoforms, AtIPK2α and AtIPK2β. Notably, the activity of these IPMK proteins is critical for heat stress acclimation in *Arabidopsis*. During heat stress, the expression of genes encoding various heat shock proteins controlled by the heat shock factors (HSFs) is affected in the AtIPK2-deficient plants. Furthermore, we show that the transcription activity of HSF is regulated by the AtIPK2 proteins. Our parallel investigations using the liverwort *Marchantia polymorpha* suggest that the InsP_6_ kinase activity of IPMK and the role of IPMK in regulating the heat stress response are evolutionarily conserved. Collectively, our study indicates that IPMK has played a critical role in transducing environmental cues for biological processes during land plant evolution.

## Introduction

Inositol phosphates (InsP) are phosphoesters of *myo*-inositol (Ins) that are synthesized by the combinatorial phosphorylation of hydroxyl group (-OH) of *myo*-inositol ring forming InsP_1_-InsP_6_. Diphosphate-containing inositol phosphates, also known as inositol pyrophosphates (PP-InsPs) are sub-family of InsP and are versatile metabolites characterized by the presence of ‘‘high energy’’ phosphoanhydride bonds, with InsP_7_ and InsP_8_ being the most characterized species (Barker et al., 2009; Chabert et al., 2023; Monserrate and York, 2010; Nguyen Trung et al., 2022; Shears, 2015; Thota and Bhandari, 2015; Wilson et al., 2013). In yeast and metazoans, PP-InsPs serve as the critical cellular messengers controlling a large array of physiological processes including phosphate homeostasis (Chabert et al., 2023; Lee et al., 2007; Li et al., 2020; Wilson et al., 2019), cellular energetics (Szijgyarto et al., 2011) and metabolism (Gu et al., 2021; Qin et al., 2023). To date, the metabolic pathways leading to the production of PP-InsPs are well established in yeast, amoeba and metazoans (Desfougeres et al., 2022; Mulugu et al., 2007; Saiardi et al., 1999; Wang et al., 2011). With the ease of genetic manipulation, the budding yeast (*Saccharomyces cerevisiae*) served as a unique model system to study the InsP metabolism (Odom et al., 2000; Wilson et al., 2013). The mammalian IP6K/yeast Kcs1-type proteins phosphorylates the C5 position of InsP_6_ and 1-InsP_7_ to generate 5-InsP_7_, and 1,5-InsP_8_, respectively (Draskovic et al., 2008; Saiardi et al., 1999). Furthermore, mammalian PPIP5K/yeast Vip1 kinases phosphorylate the C1 position of InsP_6_ and 5-InsP_7_ to generate 1-InsP_7_ and 1,5-InsP_8_, respectively (Dollins et al., 2020; Lin et al., 2009; Mulugu et al., 2007; Wang et al., 2011).

Notably, the metabolic pathway of PP-InsP production is partially conserved in plants (Cridland and Gillaspy, 2020; Laha et al., 2021b; Lorenzo-Orts et al., 2020; Riemer et al., 2022). For instance, plants lack the canonical Kcs1/IP6K-type proteins responsible for 5-InsP_7_ production found in yeast and metazoans. By contrast, *Arabidopsis* ITPK1 and ITPK2 that are not sequence related to yeast Kcs1 or mammalian IP6K enzymes, phosphorylates InsP_6_ to generate 5-InsP_7_ *in vitro* (Laha et al., 2019; Olusegun Adepoju; Whitfield et al., 2020; Zong et al., 2022) and *in planta* (Laha et al., 2022; Riemer et al., 2021). Notably, Vip1 isoforms could be identified in all available plant genomes (Desai et al., 2014; Laha et al., 2015). *Arabidopsis* Vip1 isoforms, VIH1 and VIH2 contribute to 1-InsP_7_ and InsP_8_ synthesis *in planta* (Laha et al., 2015; Riemer et al., 2021; Zhu et al., 2019). Collectively, the current literatures highlight the existence of an intricate PP-InsP metabolism and signalling network in the plant kingdom.

Thanks to these recent advances in understanding PP-InsP metabolism in plants that have created a platform to investigate the physiological functions of various PP-InsP species in plants. For instance, reduction in InsP_8_ level through perturbance of VIH2 function in Arabidopsis leads to the compromised immune response against insect herbivores and necrotrophic fungi (Laha et al., 2015; Laha et al., 2016). Furthermore, VIH-derived PP-InsPs play crucial role in regulating phosphate starvation response (Dong et al., 2019; Zhu et al., 2019). The Chlamydomonas Vip1 controls nutrient sensing (Couso et al., 2016). Similarly, ITPK1 function is also implicated in phosphate homeostasis in Arabidopsis (Kuo et al., 2018; Riemer et al., 2021). Both ITPK1 and ITPK2 regulates auxin-dependent physiological processes(Laha et al., 2022). Enzymes responsible of PP-InsP production are also linked with plant immunity against pathogenic bacteria in Arabidopsis (Gulabani et al., 2022). Collectively, these studies reinforce that PP-InsPs are critical cellular messengers regulating various aspect of plant physiology and development.

Two independent studies have reported the presence of a new PP-InsP isomer with yet unknown enantiomeric identity, 4/6-InsP_7_ in the Arabidopsis tissue extracts (Laha et al., 2022; Riemer et al., 2021). In the social amoeba *Dictyostelium discoideum*, 4/6-InsP_7_ represents the most abundant InsP_7_ species and is synthesized by the amoeba Ip6k (Desfougeres et al., 2022). Since, plants lack the canonical IP6K, it remained completely elusive how 4/6-InsP_7_ is produced in plant cells. Furthermore, whether 4/6-InsP_7_ is specific to a certain plant lineage or is ubiquitous in land plants is yet to be explored. Consequently, physiological functions of this newly identified PP-InsP species remained completely unknown.

In this study, we demonstrate that 4/6-InsP_7_ is the most abundant form of InsP_7_ present across land plants. To identify the protein(s) responsible for 4/6-InsP_7_ production in plants, we performed a structural-based homology screening, where we identified Arabidopsis IPK2α, a member of inositol polyphosphate multikinase (IPMK) family (Dovey et al., 2018; Odom et al., 2000; Saiardi et al., 2000; Stevenson-Paulik et al., 2005; Stevenson-Paulik et al., 2002; Wang and Shears, 2017; Xia et al., 2003), as the primary hit. Using *in vitro* biochemical assays, we show that indeed AtIPK2α phosphorylates InsP_6_ to 4/6-InsP_7_ *in vitro*. The *Saccharomyces cerevisiae* IPK2 phosphorylates 3-OH and 6-OH position of inositol ring generating various inositol phosphate species that serve as precursor for InsP_6_ (Odom et al., 2000). In agreement with this, the yeast *ipk2* knockout strains do not have detectable level of InsP_6_ (Odom et al., 2000). Notably, Arabidopsis IPK2-defective seedlings did not show any significant reduction in InsP_6_ level. Moreover, analyses of IPK2-deficient plants revealed that Arabidopsis IPMK isoforms control the synthesis of 4/6-InsP_7_ *in planta*. Hence, we report a non-archetypal function of IPK2 proteins in phosphorylating InsP_6_ to generate 4/6-InsP_7_ species *in planta*. Next, we explore the physiological role of AtIPK2 isoforms and reveal that the activity of AtIPK2 proteins is critical for plant adaptation to heat stress. We further show that the IPK2 controls the transcriptional activity of HSF that regulates heat stress responses through inducing transcription of the heat stress responsive genes. Our parallel investigations using the bryophyte species *Marchantia polymorpha*, the first plant to conquer the land 500 million years ago (Bowman, 2022) allow us to conclude that the functions of IPMK in controlling cellular levels of 4/6-InsP_7_ and facilitating heat stress acclimation are conserved in land plants.

## Results

### 4/6-InsP_7_ is the major form of PP-InsP species in land plants

To investigate whether 4/6-InsP_7_ is ubiquitous across land plants, we aimed to analyze the inositol phosphate profile of the representative species in each clade of the embryophytes. To this end, inositol phosphates from the thallus of *Marchantia polymorpha* (liverworts; Bryophyta), sporophylls of *Selaginella sp.* (Lycopods; Pteridophyta) and seedlings of *Arabidopsis thaliana* (Eudicot, Angiosperms) were extracted and analyzed using the recently developed capillary electrophoresis coupled with mass spectrometry method (CE-MS)(Liu et al., 2023; Qiu et al., 2023; Qiu et al., 2020). Our analyses demonstrated the presence of 4/6-InsP_7_ in all plant extracts used in the study (Fig. 1A). Furthermore, the quantification of PP-InsP species revealed that the cellular concentration of 4/6-InsP_7_ was consistently higher than that of other InsP_7_ isomers, i.e., 5-InsP_7_ and 1/3-InsP_7_ detected in the plant tissue extracts under the studied conditions (Fig. 1A). Altogether, these results demonstrate that 4/6-InsP_7_ is ubiquitous and the most abundant InsP_7_ isomer in the embryophytes analyzed in our study. Our CE-MS further identified various InsP_3-4-5_ species that are conserved in land plants (Supplementary Fig. 1).

**Figure 1.**
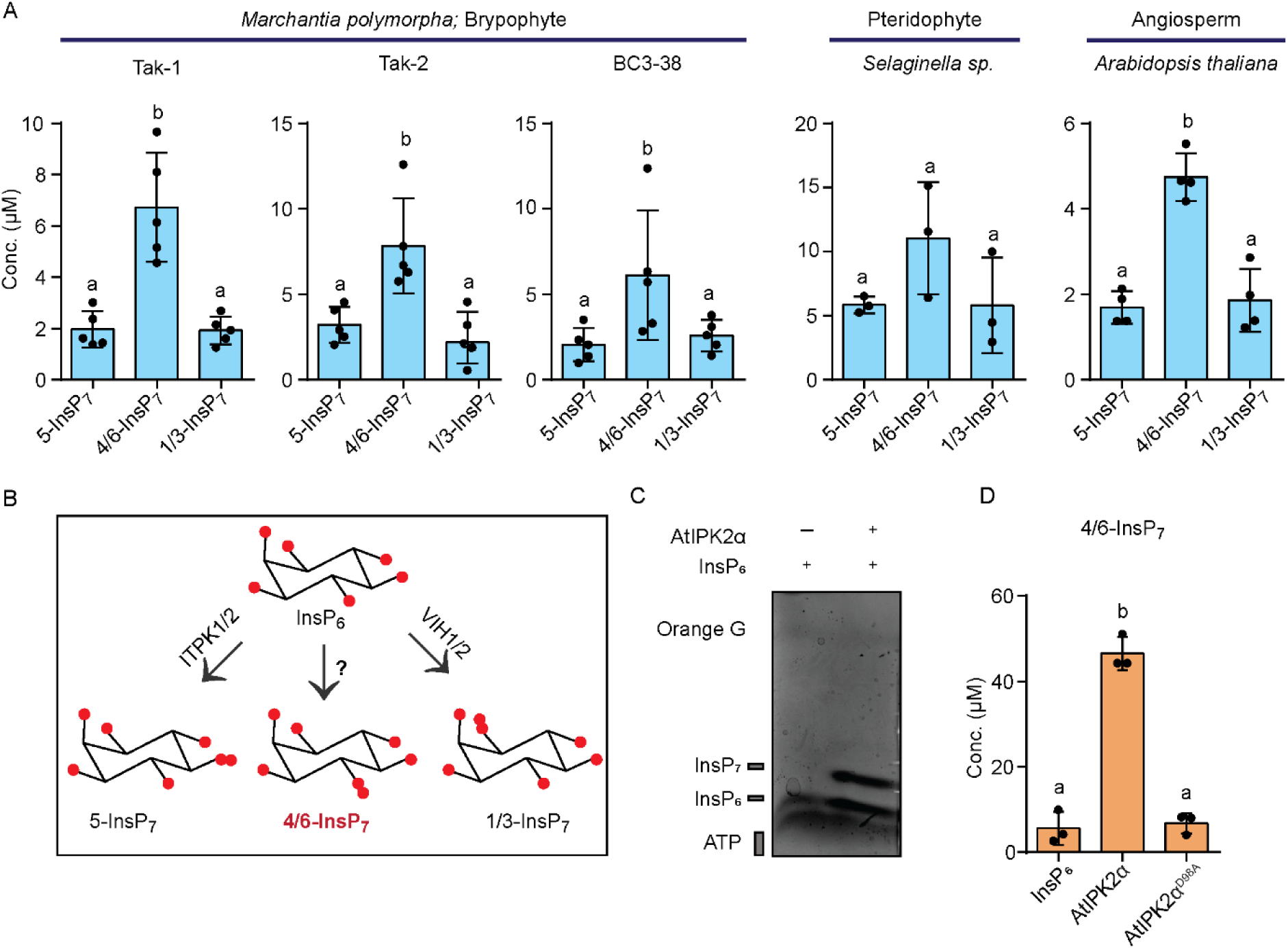
4/6-InsP_7_ represents the most abundant form of InsP_7_ isomer and is synthesized by the IPMK-type proteins *in vitro*. **A**. 4/6-InsP**_7_** is the major InsP**_7_** isomer detected in different plant extracts. CE-MS analyses of InsP**_7_** isomers present in the extracts of the designated embryophytes. 5-InsP**_7_**, 4/6-InsP**_7_** and 1/3-InsP**_7_** were assigned by mass spectrometry and identical migration time compared with their heavy isotopic standards. Data are means ± SE (n ≥ 3, biological replicates). Different letters indicate significance in one-way analysis of variance (ANOVA) followed by Tukey’s test (a and b, *P* <0.001, for Tak-1 and Tak-2, *P*<0.05 for BC3-38 and *P*<0.0001 for Col-0). **B**. Schematic representation of PP-InsP metabolism in *Arabidopsis thaliana*. InsP**_6_** is phosphorylated at position C5 by ITPK1/2 to form 5-InsP**_7_** (Laha et al., 2019; Olusegun Adepoju; Whitfield et al., 2020; Zong et al., 2022). VIH1/2 phosphorylate InsP**_6_** at position C1 to produce 1/3-InsP**_7_** (Dong et al., 2019; Riemer et al., 2021; Zhu et al., 2019). However, the protein(s) responsible for 4/6-InsP**_7_** synthesis remained unidentified. **C**. PAGE analysis of the *in vitro* kinase reaction products of AtIPK2α. The reaction product was separated by 33 % PAGE and visualized with toluidine blue. **D**. AtIPK2α synthesizes 4/6-InsP**_7_** *in vitro*. CE-MS analyses of AtIPK2α-derived *in vitro* reaction products. Data represent means ± SE (n = 3, replicates). Different letters indicate significance in one-way analysis of variance (ANOVA) followed by Dunnett’s test (a and b, *P* <0.0001).

### AtIPK2α phosphorylates InsP_6_ to form 4/6-InsP_7_ *in vitro*

Next, we aimed to identify the protein(s) responsible for 4/6-InsP_7_ production in plants. Given that green plants lack canonical IP6K-type proteins as found e.g., in mammals (Laha et al., 2019; Olusegun Adepoju; Saiardi et al., 1999; Whitfield et al., 2020; Zong et al., 2022), we speculated that the plant genome might encode a novel kinase that generates 4/6-InsP_7_ from InsP_6_. To this extent, we performed a homology search by Phyre^2^ software (Kelley et al., 2015) using human inositol hexakisphosphate kinase 3 (HsIP6K3) that converts InsP_6_ to 5-InsP_7_ (Draskovic et al., 2008; Saiardi et al., 1999), as a query sequence to identify proteins having similar structural fold albeit no sequence homology. Notably, the *Arabidopsis thaliana* inositol polyphosphate kinase alpha (AtIPK2α) was found to be one of the primary hits (Supplementary Data Set 2) that belongs to the inositol polyphosphate multikinase (IPMK) family and are able to phosphorylate different lower InsP species (Dovey et al., 2018; Odom et al., 2000; Saiardi et al., 2000; Stevenson-Paulik et al., 2005; Stevenson-Paulik et al., 2002; Wang and Shears, 2017; Xia et al., 2003). However, the role of IPMK in catalyzing the formation of phosphoanhydride was previously unknown. The recent identification of an IP6K that belongs to the IPMK family, responsible for the synthesis of 4/6-InsP_7_ in the social amoeba *Dictyostelium discoideum* (Desfougeres et al., 2022), further encouraged us to explore the potential of IPMK proteins to function as InsP_6_ kinase in plants. The *Arabidopsis* genome encodes two IPMK homologs, AtIPK2α and AtIPK2β (Stevenson-Paulik et al., 2002; Xia et al., 2003; Zhan et al., 2015).

To test the putative InsP_6_ kinase activity of *Arabidopsis* IPMK proteins, translational fusion of these proteins with an N-terminal histidine tag followed by a maltose binding protein (MBP) were expressed in a bacterial system (Supplementary Fig. 2A and 2B). Although AtIPK2β did not show any IP6K activity *in vitro*, incubation of recombinant AtIPK2α with InsP_6_, resulted a clear band that migrates slower than InsP_6_ as resolved by highly concentrated polyacrylamide gel electrophoresis (PAGE)(Pisani et al., 2014) (Supplementary Fig. 2C and Fig 1C). Furthermore, the molecular identity of the reaction product was elucidated by CE-MS analyses, wherein the AtIPK2α-derived reaction products were spiked with corresponding heavy stable isotope labelled standards (SIL) [^13^C_6_]1-InsP_7_, [^13^C_6_]5-InsP_7_ (Harmel et al., 2019) and [^18^O_2_]4-InsP_7_ (Haas et al., 2022). Notably, the AtIPK2α reaction product exactly comigrated with [^18^O_2_]4-InsP_7_ and additionally had the exact molecular mass of InsP_7_, showing that AtIPK2α is the kinase responsible for 4/6-InsP_7_ synthesis *in vitro* (Fig. 1D). Its enantiomeric identity, i.e., whether it is 4-InsP_7_ or 6-InsP_7_ cannot be dissected using this method (Ritter et al., 2023). Further, to elucidate the role of the conserved PXXXDXKXG motif (Stevenson-Paulik et al., 2002) of AtIPK2α in its catalytic activity, translational fusion of the catalytic dead mutants of AtIPK2α i.e., AtIPK2α^D98A^ and AtIPK2α^K100A^ were generated and expressed in bacteria (Supplementary Fig. 2B). Incubation of these catalytic dead mutants with InsP_6_ and ATP did not result in more polar species than InsP_6_ as determined by PAGE and CE-MS (Supplementary Fig. 2D and Fig. 1D), highlighting the importance of the key residues D98 and K100 (Fig. 1D and Supplementary Fig. 2D). A small amount of 1/3-InsP_7_ was also detected in the AtIPK2α-catalyzed reaction products (Supplementary Fig. 2E). Collectively, our analyses identified AtIPK2α as an InsP_6_ kinase that phosphorylates InsP_6_ at the C4 or C6 phosphate synthesizing 4/6-InsP_7_ *in vitro*.

### AtIPK2-type proteins are responsible for maintaining the cellular level of 4/6-InsP_7_

To investigate the contribution of the IPMK homologs to PP-InsP homeostasis *in planta*, we aimed to generate *Arabidopsis* lines with an altered expression of *AtIPK2α* or *AtIPK2β*. Since both isoforms show redundant expression in different tissues (Stevenson-Paulik et al., 2002; Xia et al., 2003) and the *atipk2βatipk2α* double knockout plants are embryonic lethal(Zhan et al., 2015), we aimed to generate knockdown lines of *AtIPK2α* in the *atipk2β* knockout plants. We established stable lines for dexamethasone-inducible RNAi gene silencing of *AtIPK2α* in the *atipk2β* knockout plants by introduction of the complete coding DNA sequence (∼900 bp) of *AtIPK2α* into the Hellsgate12 hairpin cassette under the dexamethasone-inducible bidirectional pOp6 promoter of the pOpOFF2(Hyg) vector (Parvin et al., 2017; Wielopolska et al., 2005) (Supplementary Fig. 3A). Selected independent knockdown (*kd*) lines were confirmed by genotyping (Supplementary Fig. 3B) and were tested for RNAi induction upon treatment with Dex by means of a β-glucuronidase (GUS) reporter gene under control of the bidirectional pOp6 promoter (Supplementary Fig. 3A and 3C). To explore the contribution of the two AtIPK2 isoforms in 4/6-InsP_7_ synthesis *in planta*, we studied the extracts of Col-0 (wild-type) and the *atipk2β^-/-^atipk2α^kd^*seedlings by CE-MS analyses (" ref-type="fig">Fig. 2A and 2E). The knockdown lines were indeed significantly impaired in 4/6-InsP_7_ synthesis. The level of 4/6-InsP_7_ was reduced by 40% in the independent knockdown lines compared to Col-0 plants (" ref-type="fig">Fig. 2A and 2B). In contrast, the level of InsP_6_ and other InsP_7_ isomers i.e., 5-InsP_7_ and 1/3-InsP_7_ were not altered significantly in the *atipk2β^-/-^atipk2α^kd^* seedlings (Fig. 2E). Moreover, the knockdown lines show an increased level of InsP_4_ species with yet unknown isomeric identity (Fig. 2E). Given that transcripts of the *AtIPK2* isoforms could be altered during heat shock (Kilian et al., 2007) (Supplementary Data Set 3), we asked whether heat stress influences inositol phosphate profile of the *atipk2β^-/-^atipk2α^kd^*lines compared to Col-0 plants.

**Figure 2.**
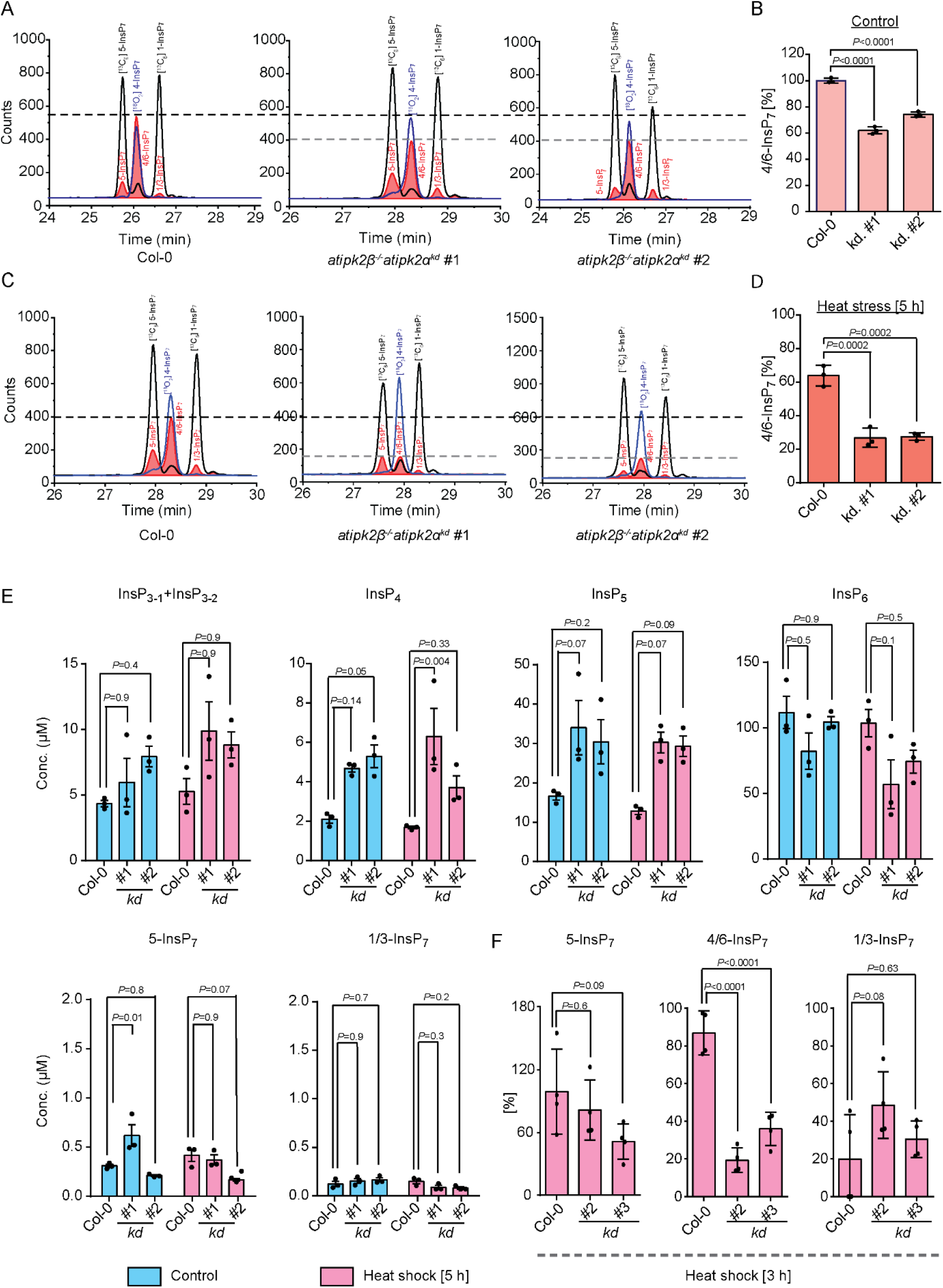
Arabidopsis AtIPK2α and AtIPK2β regulate cellular level of 4/6-InsP_7_. **A**. CE-MS analysis (extracted ion electropherograms) of InsP**_7_** in Col-0 and *atipk2β^-/-^atipk2α^kd^* line (red trace) with spiked [^13^C**_6_**] (black plot) and [^18^O**_2_**] (blue plot) labelled InsP**_7_**. 4/6-InsP**_7_** was assigned by mass spectrometry and identical migration time compared with its heavy isotopic standards ([^18^O**_2_**] 4-InsP**_7_**). **B**. Quantification of 4/6-InsP**_7_** in Col-0 and *atipk2β^-/-^atipk2α^kd^* line. 4/6-InsP**_7_** is presented as percentage to total InsP**_6_**. Data are means ± SE (n = 3, biological replicates) and significant differences were analyzed one-way analysis of variance (ANOVA) followed by Dunnett’s test. **C**. Activity of Arabidopsis IPK2 proteins is critical for maintaining 4/6-InsP**_7_** level during heat shock. CE-MS analysis (extracted ion electropherograms) of InsP**_7_** in Col-0 and *atipk2β^-/-^atipk2α^kd^*line after heat shock of 5 h at 37⁰C with spiked [^13^C**_6_**] (black plot) and [^18^O**_2_**] (blue plot) labelled InsP**_7_**. 4/6-InsP**_7_** was assigned by mass spectrometry and identical migration time compared with its heavy isotopic standards([^18^O**_2_**] 4-InsP**_7_**). **D**. Quantification of 4/6-InsP**_7_** in Col-0 and *atipk2β^-/-^atipk2α^kd^* line after heat shock for 5 h at 37⁰C. 4/6-InsP_7_ is presented as percentage to total InsP_6_. Data are means ± SE (n = 3, biological replicates) and significant differences were determined by one-way analysis of variance (ANOVA) followed by Dunnett’s test. **E**. CE-MS analyses of inositol polyphosphate levels of 2-week-old Col-0 and *atipk2β^-/-^ atipk2α^kd^* seedlings. Plants were grown on solidified half strength MS media, supplemented with 1% (w/v) sucrose, growth conditions are detailed in the methods section. Seedlings were subjected to heat shock at 37⁰C for 5 h and are harvested. Inositol phosphates were extracted by TiO2 pull down and were subjected to CE-MS analyses. The InsP_5_, InsP_6_ and InsP_7_ species were assigned by mass spectrometry and identical migration time compared with their relative standards. One InsP_4_ and two InsP_3_ isomers were detected. Data are means ± SE (n =3, biological replicates). Statistical significance is determined in two-way ANOVA followed by Tukey’s test. **F**. CE-MS analyses of different isomers of InsP_7_ in 2-week-old Col-0 and *atipk2β^-/-^ atipk2α^kd^* seedlings after heat shock of 3 h at 37⁰C. Graph represents the fold difference in InsP_7_ isomers upon heat stress in the designated genotypes. Values are ± SE (n =4, biological replicates). Statistical significance is determined in one-way ANOVA followed by Dunnett’s test.

When the plant lines were exposed to heat stress at 37⁰C for 5 h, a 40% decrease of 4/6-InsP_7_ species in the Col-0 extracts was observed (" ref-type="fig">Fig. 2A-2D). Strikingly, the *atipk2β^-/-^atipk2α^kd^* lines suffered severely in 4/6-InsP_7_ production with an approximate 60% reduction compared to the Col-0 plants when exposed heat stress for 5 h (" ref-type="fig">Fig. 2C and 2D), providing additional evidence that AtIPK2 proteins control the cellular 4/6-InsP_7_ level. Notably, we did not observe any specific changes in other InsP species including the PP-InsP isomers, 5-InsP_7_ and 4/6-InsP_7_ during heat stress (Fig. 2E). Furthermore, significant reduction of 4/6-InsP_7_ species in the *atipk2β^-/-^atipk2α^kd^* lines compared to Col-0 plants could be observed after 3 h of heat shock treatment (Fig. 2F). Collectively, our data suggest that *Arabidopsis* IPMK isoforms are critical for 4/6-InsP_7_ production *in planta.* The remaining pool of 4/6-InsP_7_ species in the *atipk2β^-/-^ atipk2α^kd^*lines might originate from the residual activity of AtIPK2α in the knockdown lines.

### AtIPK2α and AtIPK2β cooperate to control heat stress acclimation in *Arabidopsis*

To elucidate the possible role of AtIPK2α and AtIPK2β in heat stress acclimation, we exposed Col-0, both the single knockout lines and the *atipk2β^-/-^atipk2α^kd^* lines to various temperatures and monitored the adaptive responses commonly referred as thermomorphogenesis (Casal and Balasubramanian, 2019; Delker et al., 2022; Quint et al., 2023). Under the control condition (22⁰C), all genotypes including Col-0, *atipk2β*, *atipk2α* and the knockdown lines did not exhibit any obvious growth difference including primary root growth (" ref-type="fig">Fig. 3A and 3B). However, at higher ambient temperature (28⁰C), Col-0, *atipk2β* and *atipk2α* displayed longer primary root length (" ref-type="fig">Fig. 3A and 3B). In contrast, the temperature-induced root elongation was completely attenuated in the independent knockdown lines (" ref-type="fig">Fig. 3A and 3B), suggesting that AtIPK2α and AtIPK2β act collaboratively to regulate root thermomorphogenesis. To further interrogate the role of AtIPK2 isoforms in plant thermotolerance, we performed basal thermotolerance assays using Col-0, *atipk2β*, *atipk2α* and the three independent knockdown lines (" ref-type="fig">Fig. 3C-3E and Supplementary Fig. 4B). Seedlings were subjected to heat stress as indicated in the method section (Fig. 3D) and plant survival was assessed, defined by their ability to maintain and generate fresh green leaves (Fig. 3C and Supplementary Fig. 4B). Notably, we found that the *atipk2β^-/-^atipk2α^kd^* lines were severely affected by heat stress showing poor survival rate and thus exhibited severely compromised thermotolerance as compared to Col-0 and the single mutant lines (" ref-type="fig">Fig. 3C and 3E). This highlights the role of AtIPK2 isoforms in maintaining a plant’s basal thermotolerance ability. We also examined the shoot response after exposing all the genotypes at the high ambient temperature. Similar to root thermomorphogenesis, the *atipk2β^-/-^atipk2α^kd^* lines showed compromised hypocotyl elongation when exposed to elevated temperature (" ref-type="fig">Fig. 3F and 3G) indicating that AtIPK2 contributes to shoot adaptive response to heat stress as well. In conclusion, our data suggest that AtIPK2α and AtIPK2β function redundantly to regulate heat stress acclimation in *Arabidopsis*.

**Figure 3.**
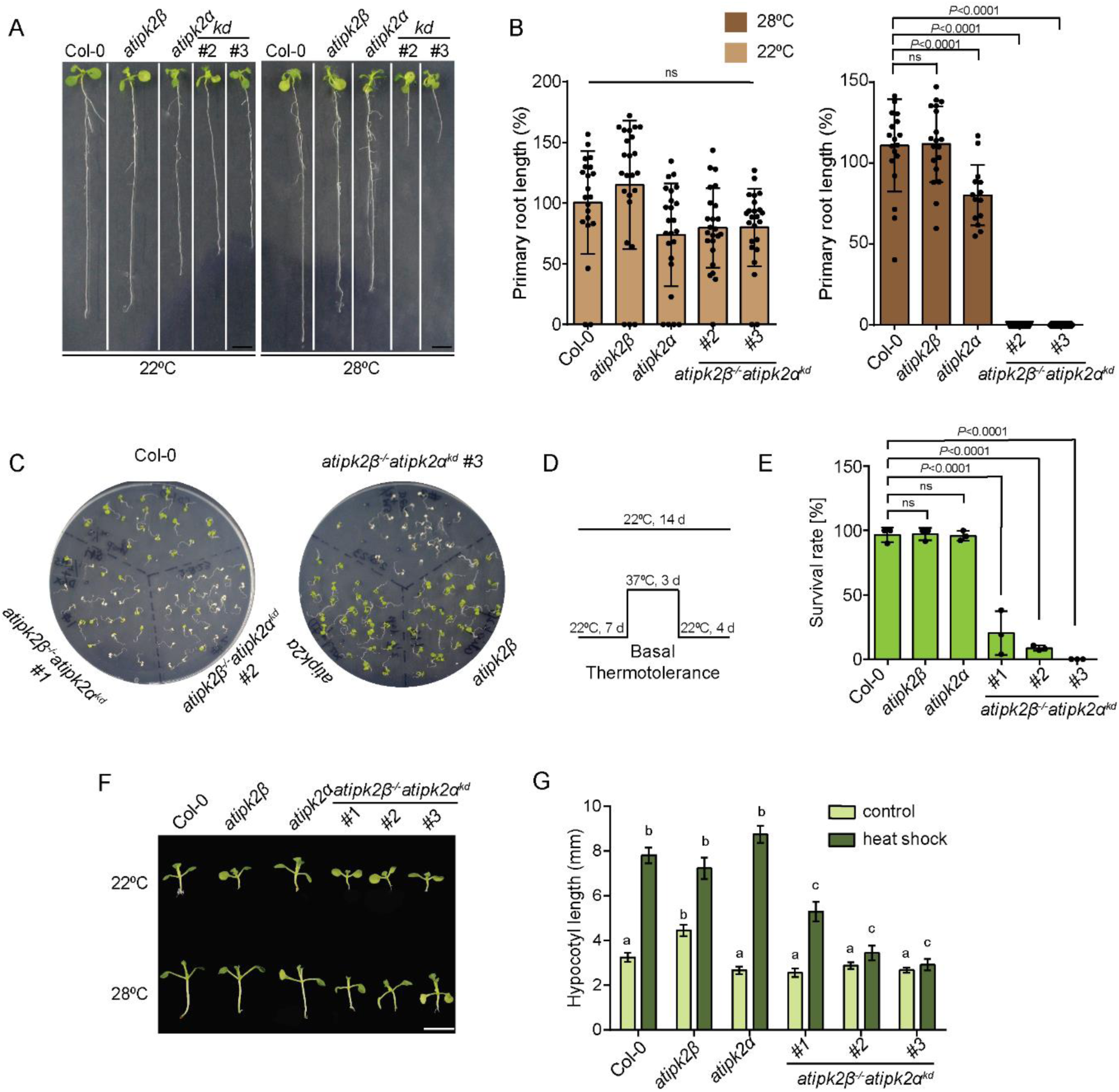
AtIPK2α and AtIPK2β control heat stress acclimation in *Arabidopsis*. **A.** AtIPK2α and AtIPK2β control root thermomorphogenesis. Photograph showing root thermomorphogenesis phenotype of Col-0, *atipk2β, atipk2α* and *atipk2β^-/-^atipk2α^kd^* lines (*kd*). 7-day-old seedlings were exposed to 28°C for 5 days. Control plates were maintained at 22⁰C. Images were taken after 5 days of heat shock along with the non-treated plants. Scale bar = 5 mm. **B.** Quantification of primary root length of Col-0, *atipk2β, atipk2α* and *atipk2β^-/-^atipk2α^kd^* lines grown at 22⁰C and 28⁰C. Root lengths were evaluated by ImageJ software. Values are ±SE, (n ≥30) and significant differences were determined by one-way analysis of variance (ANOVA) followed by Dunnett’s test. **C.** Activity of Arabidopsis IPMK isoforms is critical for survival during heat stress. Photograph showing basal thermotolerance phenotype of Col-0, *atipk2β, atipk2α* and *atipk2β ^-/-^atipk2α^kd^* lines after heat stress at 37⁰C. Surviving seedlings maintain green leaves and show emerged new leaves. **D.** Simplified experimental setup used for analysis of the survival phenotype. 7-day-old plants were exposed to 37⁰C for 3 days and were subjected to subsequent recovery at 22⁰C for 4 days. Control plates were maintained at 22⁰C. **E.** Survival rate analysis of the designated genotypes at 37⁰C. Values are means ±SE, (n=3, biological replicates) with each data point indicated and significant difference is determined by one-way analysis of variance (ANOVA) followed by Dunnett’s test. The experiment was repeated several times with independent generation. **F.** AtIPK2α and AtIPK2β regulate hypocotyl elongation during heat stress. Representative photograph of high temperature-induced hypocotyl elongation phenotype of Col-0, *atipk2β, atipk2α* and *atipk2β ^-/-^ atipk2α^kd^* lines grown at 22⁰C and 28⁰C. 7-day-old seedlings were exposed to 28°C for 5 days. Control plates were maintained at 22⁰C. Images were captured after 5 days of heat stress. Scale bar = 6mm **G.** Quantification of high temperature-induced hypocotyl elongation of the designated genotypes grown at 22⁰C and 28⁰C. Hypocotyl elongation was evaluated by using ImageJ. Data are means ± SE (n ≥20, biological replicates). Different letters indicate significance in two-way analysis of variance (ANOVA) followed by Tukey’s test (a and b, *P*<0.0001; a to c, *P*<0.0001).

### IPK2 homologs are ubiquitous across plant kingdom and InsP_6_ kinase activity of IPK2 is conserved in the liverwort *M. polymorpha*

Next, to explore the functional conservation of InsP_6_ kinase activity of AtIPK2 proteins in land plants, we constructed a phylogenetic tree including diverse taxa of plant kingdom such as green algae (Chlorophyta), mosses (Bryophyta), lycopods (Pteridophyta), monocot and eudicot (Angiosperms) (Supplementary Data Set 5). The phylogenetic analysis allowed us to identify genes encoding IPMK-type proteins across the plant kingdom (Supplementary Fig. 5A). The analyses further suggest that these IPMK/IPK2 homologs are derived from a single ancestral gene, with subsequent radiation in the individual lineages (Supplementary Fig. 5A). To understand the ancestral function of plant IPK2α-type kinases in inositol phosphate homeostasis, we began to characterize the homolog of AtIPK2 in the liverwort *M. polymorpha*, a bryophyte whose genome sequence is available and is emerging as a model plant species to study land plant evolution (Bowman et al., 2017; Kohchi et al., 2021). The *M. polymorpha* genome encodes a single IPK2α homologue named MpIPMK (as per nomenclature guidelines of *Marchantia*) (Bowman et al., 2016) (Supplementary Fig. 5A).

The multiple sequence alignment of MpIPMK with AtIPK2α and yeast Ipk2 showed that MpIPMK possesses the conserved residues of the PXXXDXKXG catalytic motif suggesting that MpIPMK could be a functional homolog of AtIPK2 proteins (Supplementary Fig. 5B and 6A). Comparison of structural model of MpIPMK with AtIPK2α (Supplementary Fig. 6B-6D) further indicates that MpIPMK could function as AtIPK2α. We validated this hypothesis by taking advantage of the yeast (*Saccharomyces cerevisiae*) *ipk2*Δ knockout strain and studied the consequences of heterologous expression of Mp*IPMK* in these mutant strains. The *S. cerevisiae* genome encodes a single IPMK homolog Ipk2(Odom et al., 2000; Saiardi et al., 2000). Ectopic expression of Mp*IPMK* could restore the *ipk2*Δ-associated growth defects at high temperature (37⁰C) (Fig. 4B). Rescue of the *ipk2*Δ-associated growth defect could only be noticed by the ectopic expression of wild-type MpIPMK but not by the catalytic dead mutant MpIPMK^D130A^, MpIPMK^K132A^ and MpIPMK^D130AK132A^, indicating that MpIPMK possesses similar catalytic activity to yeast Ipk2 and AtIPK2α/AtIPK2β (Fig. 4B). This conclusion was further substantiated by the HPLC analyses of the yeast *ipk2*Δ transformants expressing Mp*IPMK* and Mp*IPMK^D130AK132A^* (Fig. 4C). The altered InsP profile of the yeast *ipk2*Δ strain could be fully rescued by the ectopic expression of Mp*IPMK* (Fig. 4C). In contrast, the catalytically dead variant of MpIPMK, MpIPMK^D130AK132A^ fails to rescue the defective InsP profile (Fig. 4C). These data allowed us to conclude that MpIPMK is a functional yeast Ipk2 homolog encoded by the *M. polymorpha* genome. To delineate whether MpIPMK possesses InsP_6_ kinase activity similar to AtIPK2α, we incubated the translational fusion polypeptides of MpIPMK along with its catalytic dead variants (MpIPMK^D130A^, MpIPMK ^K132A^, MpIPMK ^D130AK132A^) with InsP_6_ and ATP and the reaction products were resolved by PAGE (Supplementary Fig 6E,6F and Fig. 4D). The PAGE analyses revealed the presence of a clear kinase product when InsP_6_ was incubated with the wild-type IPMK protein (Fig. 4D) suggesting that MpIPMK can phosphorylate InsP_6_ *in vitro*. The mutant MpIPMKs failed to synthesize more polar species of InsP_6_ (Fig. 4D). To get further insights into MpIPMK catalytic activity, we expressed Mp*IPMK* in different mutant yeast strains defective in PP-InsP metabolism. The expression of Mp*IPMK* did not rescue the *kcs1*Δ-associated growth defect (Supplementary Fig. 7A). Similarly, MpIPMK was not able to rescue the *vip1Δ*-associated growth defects (Supplementary Fig. 7B). Collectively, these data indicate that MpIPMK is a functional yeast Ipk2 homolog and does not possess either Kcs1-type or Vip1-type catalytic activity. To further corroborate the catalytic activity of MpIPMK and to deduce the structural identity of the MpIPMK reaction product, we subjected the *in vitro* MpIPMK-derivatives for CE-MS analyses. Notably, the MpIPMK reaction product showed exact comigration with the [^18^O_2_]4-InsP_7_ standard, demonstrating that MpIPMK is the kinase responsible for 4/6-InsP_7_ synthesis *in vitro* (Fig. 4D). Collectively, all these data unveil that MpIPMK is a novel InsP_6_ kinase responsible for the synthesis of 4/6-InsP_7_ *in vitro*. Similar to the AtIPK2 reaction products, a small amount of 1/3-InsP_7_ could be detected using CE-MS analyses in the MpIPMK-derived reaction products (Supplementary Fig. 7C).

**Figure 4.**
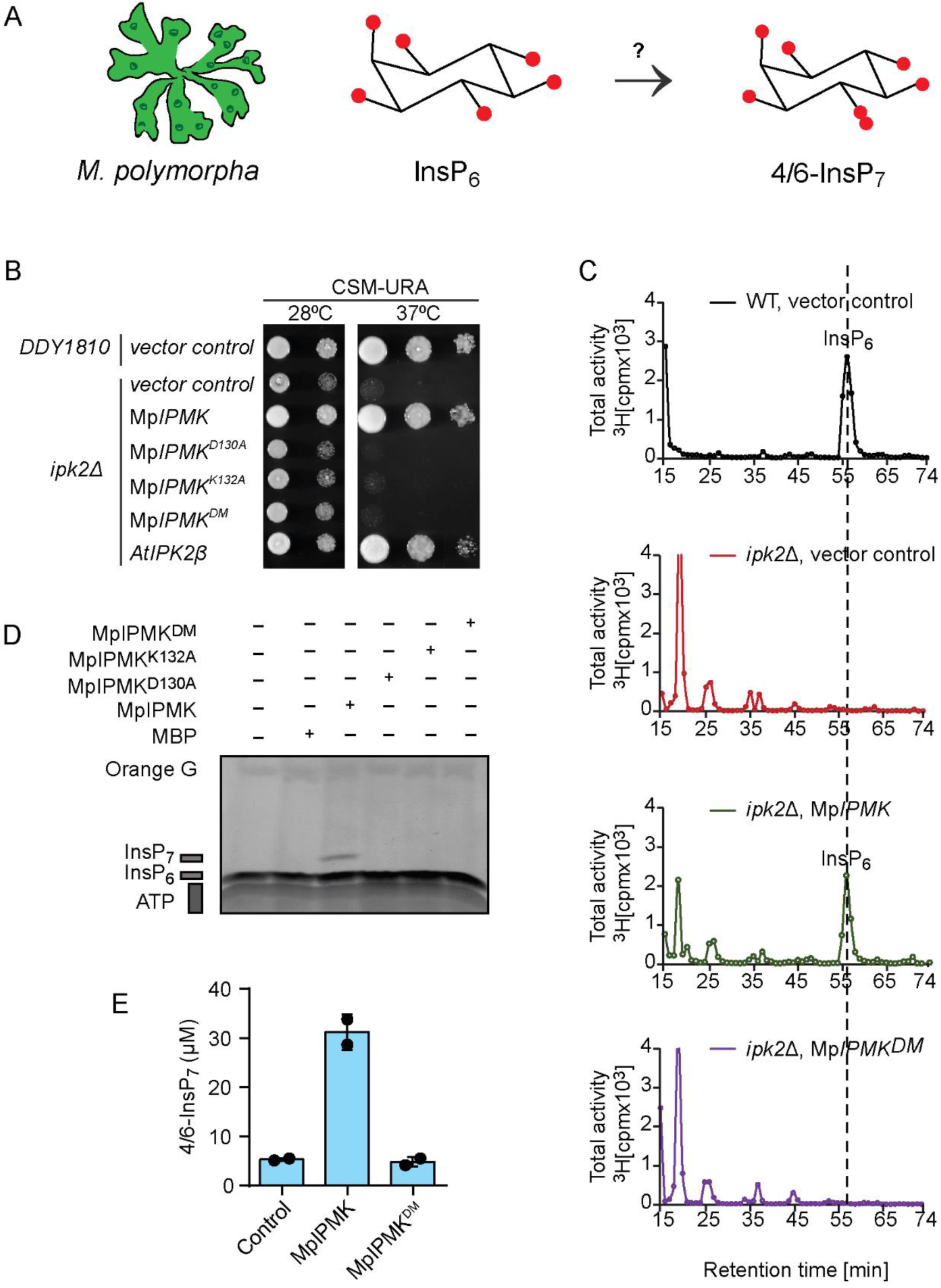
*M. polymorpha* genome encodes a functional yeast Ipk2 homolog, MpIPMK that generates 4/6-InsP_7_ *in vitro*. **A**. Schematic representation of 4/6-InsP_7_ synthesis in *M. polymorpha*. **B**. MpIPMK is a functional yeast Ipk2 homolog. Complementation of *ipk2*Δ-associated growth defects in yeast by ectopic expression of Mp*IPMK*, the AtIPK2 homolog in *M. polymorpha*. The *ipk2Δ* yeast strain transformed with episomal pCA45 (*URA3*) plasmid carrying Mp*IPMK* with N-terminal GST translation fusion were spotted in 8-fold serial dilution onto uracil-free plate and were incubated at 28°C and 37⁰C for 3 days. *AtIPK2β* served as positive control (Stevenson-Paulik et al., 2002; Xia et al., 2003) and empty vector served as negative control. **C**. Complementation of the altered InsP profile of *ipk2Δ* yeast mutant by ectopic expression of Mp*IPMK*. HPLC profiles of extracts from [^3^H] inositol-labelled yeast transformants. Extracts were resolved by SAX-HPLC, and fractions collected each minute for subsequent determination of radioactivity as indicated. Experiments were repeated two times with similar results. **D**. MpIPMK phosphorylates InsP_6_ *in vitro*. Recombinant His_8_-MBP-MpIPMK and the catalytic dead proteins were incubated with 12.5 mM ATP, and 10 nmol InsP_6_ at 37⁰C for 12 h in reaction buffer. The reaction product was separated by 33 % PAGE and visualized with toluidine blue. Experiments were repeated independent times with similar results. **E**. Production of 4/6-InsP_7_ by MpIPMK *in vitro*. Quantification of MpIPMK-derived 4/6-InsP_7_ using CE-MS. Data represent means ± SEM (n=2, replicates). Experiments were repeated two times with similar results.

### MpIPMK contributes to 4/6-InsP_7_ synthesis *in planta* and controls heat stress responses

To decipher the contribution of MpIPMK in PP-InsP homeostasis, independent knockout lines of Mp*IPMK* were generated using CRISPR-Cas9 technology (Sugano et al., 2018; Sugano et al., 2014) in the wild-type (WT) Tak-1 *M. polymorpha* plants (Fig. 5A and Supplementary Fig. 8A). These mutant plants did not show any obvious growth defects compared to the WT plants (Supplementary Fig. 8B). To assess the consequence of altered expression of Mp*IPMK* in inositol phosphate metabolism, we performed CE –MS analyses of WT and the independent Mp*ipmk* knockout plants (" ref-type="fig">Fig. 5B-5C and Supplementary Fig. 8C). Notably, MpIPMK-defective plants are compromised significantly in their 4/6-InsP_7_ level suggesting that indeed MpIPMK contributes to 4/6-InsP_7_ production *in planta* (Fig. 5B). This analysis also indicates that the *M. polymorpha* genome encodes protein(s) other than IPMK responsible for 4/6-InsP_7_ production in thallus. Moreover, Mp*ipmk* knockout plants show increased levels of different InsP_3_ species (Supplementary Fig. 8C).

**Figure 5.**
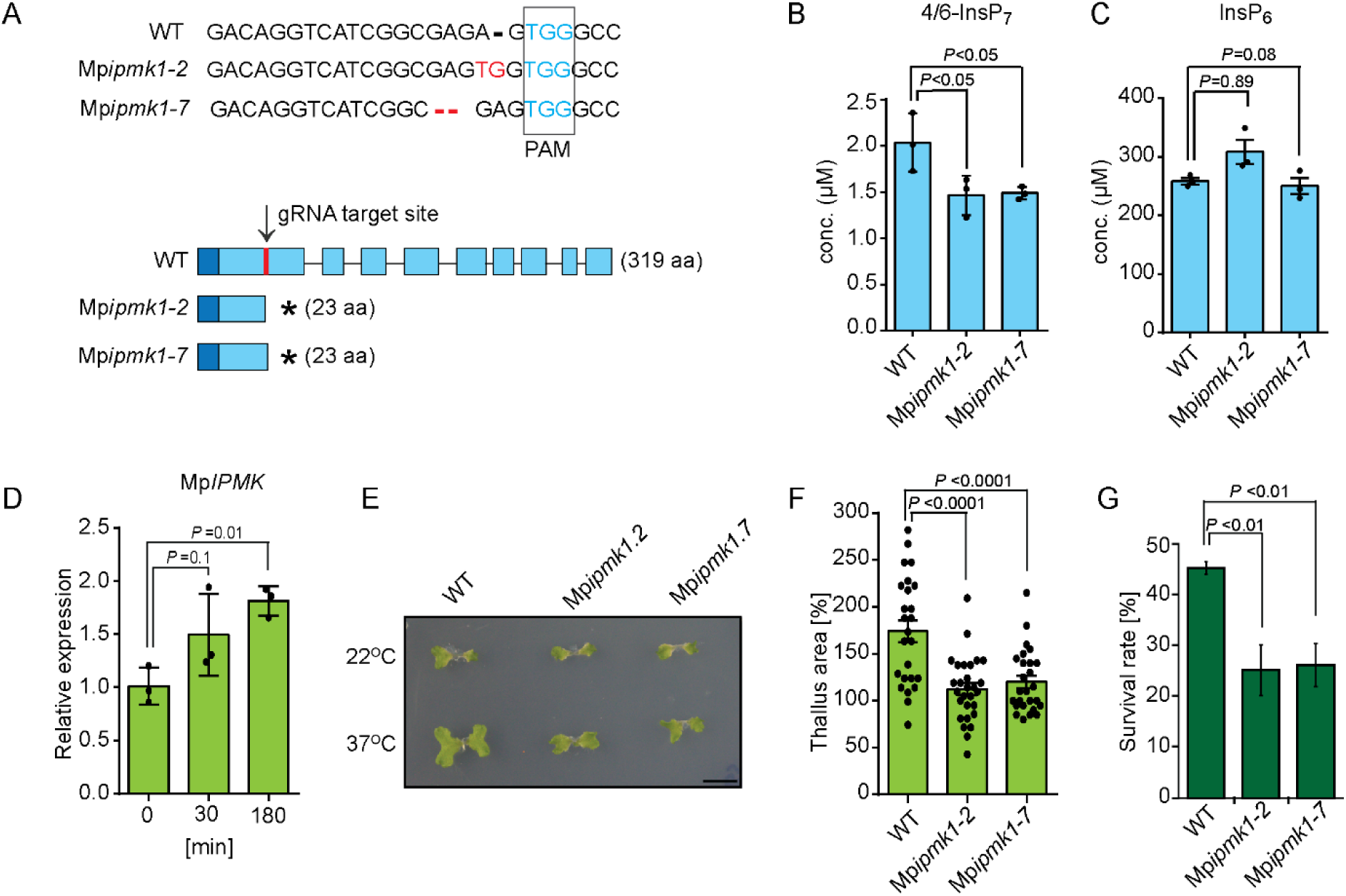
MpIPMK controls production of 4/6-InsP_7_ *in planta* and heat stress acclimation by IPMK-type protein is conserved in the early land plant, *M. polymorpha*. **A**. Generation of Mp*ipmk* knockout lines. Schematic representation of the two independent Mp*ipmk* knockout lines obtained by CRISPR-Cas9 gene editing technology. Mp*ipmk1-2* has insertion of nucleotide preceding PAM sequence while Mp*ipmk1-7* has deletion of two nucleotide preceding PAM sequence. The mutations resulted in a premature stop codon. **B**. MpIPMK-deficient plants are compromised in 4/6-InsP_7_ production. CE-MS analyses of 4/6-InsP_7_ level in 14-day-old thalli of WT, Mp*ipmk1.2* and Mp*ipmk 1.7* knockout lines. 4/6-InsP_7_ is presented in molar concentration (µM). Data are means ± SE (n = 3, biological replicates). Significant difference is determined by one-way ANOVA followed by Dunnett’s test. **C**. CE-MS analyses of InsP_6_ levels of 2-week-old WT, Mp*ipmk1.2* and Mp*ipmk1.7* knockout lines as indicated. Data are means ± SE (n = 3, biological replicates). Significant difference is determined by one-way ANOVA followed by Dunnett’s test. Experiments were repeated independent times with similar results. **D**. Quantitative RT-PCR (qRT-PCR) analysis of Mp*IPMK* expression in WT plants exposed to heat stress for 30 min and 180 min at 37⁰C. *MpACT7* is used as an internal control. **E**. MpIPMK activity is indispensable for inducing heat stress response. Photograph showing representative plants of WT and Mp*ipmk1.2* and Mp*ipmk1.7* thalli grown at 22⁰C (top panel) and plants grown at 37⁰C for 8 h in growth chamber with a subsequent recovery at 22⁰C (bottom panel) for 14 days. Control plates were maintained at 22⁰C. **F**. Quantification of thallus area as a measure of thermomorphogenetic response. 5-day-old gemmalings of WT and Mp*ipmk* knockout lines were subjected to heat stress at 37⁰C for 8 h in plant chamber followed by subsequent recovery at 22⁰C for 14 days. The experiment was repeated several times with independent generation. Values are means ±SE (n=3, biological replicates) and significant difference is determined by one-way analysis of variance (ANOVA) followed by Dunnett’s test. **G**. Quantification of survival rate representing the basal thermotolerance phenotype of WT and Mp*ipmk* knockout lines. 5-day-old gemmalings of each genotype were subjected to heat stress at 37⁰C in water bath for 8 h followed by subsequent recovery at 22⁰C for 4 days. The survival of the plant after heat stress was accessed by the ability of the plant to maintain green thallus. Values are means ± SE (n=3, biological replicates and each replicate represents ≥ 15 gemmalings). Statistical difference is determined by one-way ANOVA followed by Dunnett’s test.

Similar to the Arabidopsis knockdown seedlings, no changes in the InsP_6_ level could be observed in the Mp*ipmk* knockout thalli (Fig. 5C) suggesting that the archetypal activity of yeast IPK2 to contribute to InsP_6_ production is not conserved in the vegetative tissues of land plants.

Given the role of AtIPK2α and AtIPK2β proteins in heat stress acclimation and that Mp*IPMK* transcript level is induced after exposure to heat stress (Fig. 5D), we asked whether MpIPMK is involved in regulating thermomorphogenesis in *M. polymorpha*. To this end, we monitored the response of WT and the two independent Mp*ipmk* mutant plants after exposing them to the elevated temperature, 37⁰C. In line with previous reports (Ludwig et al., 2021; Wu et al., 2022), heat treatment resulted in increased thallus area of the WT plants (Fig. 5E). In contrast, the Mp*ipmk* mutants displayed severe reduction in thallus area compared to the WT plants suggesting that Mp*ipmk* mutants have compromised resilience to heat stress (" ref-type="fig">Fig. 5E and 5F). To corroborate further, we monitored the survival rate of WT plants and the Mp*ipmk* knockout lines after permissive heat stress and the subsequent recovery at 22⁰C for 4 days. Remarkably, the Mp*ipmk* knockout lines showed poor thermotolerance when compared with the WT plants as evident from the survival percentage of the mutant lines after heat stress (Fig. 5G). Taken together, these data show that MpIPMK critically contributes to the plant resilience to heat stress.

### IPK2 promotes transcription activity of HSF

To gain mechanistic insights, we first tested whether genes encoding members of HSP families, that play role in heat stress acclimation (Hahn et al., 2011; Li et al., 2023; Yang et al., 2022), are differentially expressed in the knockdown lines. Our analyses showed that the expression of different *HSP*s is largely compromised in the *atipk2β^-/-^atipk2α^kd^*lines when compared to Col-0 plants (" ref-type="fig">Fig. 6A and 6B). Considering that the *atipk2β^-/-^atipk2α^kd^*lines suffered with the compromised expression of genes that are regulated by different heat shock transcription factors (HSF-TFs) and that IPMK/AtIPK2 is already implicated in transcriptional regulation in various eukaryotes (Beon et al., 2022; Malabanan and Blind, 2016; Odom et al., 2000; Sang et al., 2017), we asked whether IPMK controls HSF activity. To this aim, we monitored the HSF activity by measuring luciferase activity under control of *HSP18.2* promoter element(Luo et al., 2022). Our analyses indicate that the presence of AtIPK2 significantly enhances the transcriptional activity of HSFA1V (Fig. 6C).

**Figure 6.**
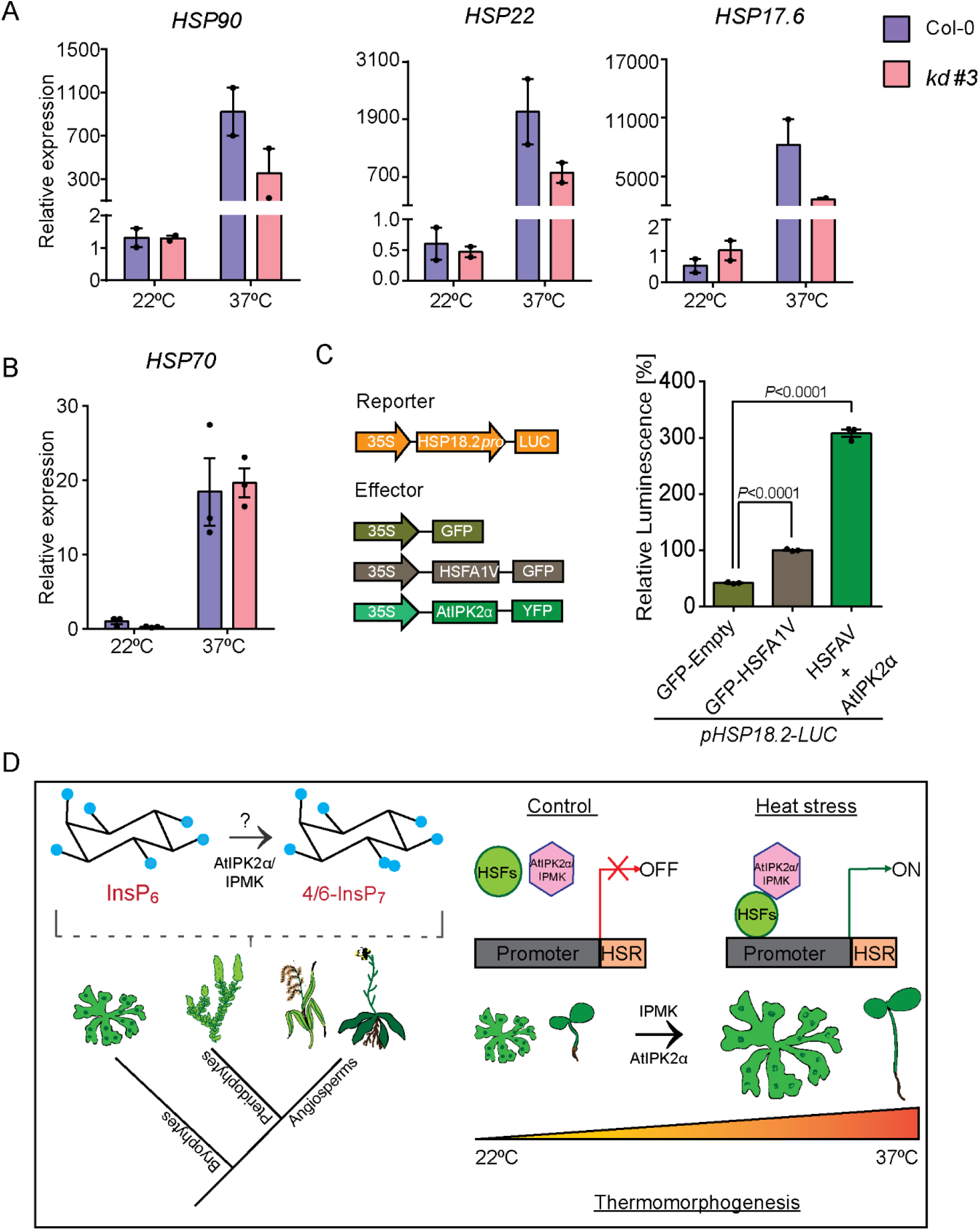
AtIPK2α modulates the transcriptional activity of HSFs. **A and B.** Quantitative RT-PCR (qRT-PCR) analysis of different *HSP*s in Col-0 and *atipk2β^-/-^atipk2α^kd^*lines after heat shock. 14-day-old seedlings were exposed to 37⁰C for 3 h and were harvested for qRT-PCR analysis. Transcript levels of the benchmark genes are presented relative *ACTIN2* transcript. Values are means ± SE (n ≤3, biological replicates). **C.** Schematic diagrams of luciferase reporter and effector constructs used in transient transactivation assays in *Nicotiana benthamiana* leaves (left panel). Statistical analysis of the expression of *pHSP18.2-LUC* in presence of HSFA1V and AtIPK2α (right panel). GFP served as control. *P* values are determined by two-way analysis of variance (ANOVA) followed by Dunnett’s test. **D.** A model illustrating the conserved function of IPMK-type protein in synthesizing 4/6-InsP_7_, the most abundant InsP_7_ species and regulating thermomorphogensis in land plants. The question mark denotes the possible existence of one or more proteins other than IPMK/AtIPK2, contributing 4/6-InsP_7_ synthesis *in planta*.

In conclusion, our study reveals that 4/6-InsP_7_ is the predominant form of PP-InsP species present in land plants. We showed that AtIPK2α generates 4/6-InsP_7_ *in vitro* and that cellular level of 4/6-InsP_7_ is controlled by AtIPK2α and AtIPK2β. Furthermore, AtIPK2 proteins act as a novel regulator for heat stress acclimation. Investigation using *M. polymorpha* allowed us to conclude that IPMK-dependent 4/6-InsP_7_ production is an ancestral function and the role of IPMK-type proteins in heat stress acclimation is evolutionarily conserved (Fig. 6D). The unaltered InsP_6_ level despite some accumulation of InsP_3_ and InsP_4_ species in Arabidopsis and *M. polymorpha* IPK2-deficient lines suggests that the archetypal activity of IPK2 in producing phytic acid in yeast (Odom et al., 2000) is not conserved in the vegetative tissue of land plants. Our data further indicates that the InsP_6_ kinase activity of IPK2 evolved during the divergence of green plants from the fungal lineage. This study also indicates that there could be one or more proteins besides IPMK involved in 4/6-InsP_7_ synthesis. Finally, our study provides mechanistic insights into the regulation of heat stress acclimation by IPK2 through modulating HSF transcription activity. How precisely IPMK controls HSF activity and therefore heat stress response are yet to be understood. Taken together, our study provides a mechanistic framework to understand the conserved role of IPMK-derived 4/6-InsP_7_ in regulating various cellular processes, and how this contributes to land plant evolution.

## Methods

### Phylogenetic analysis

The phylogenetic tree was constructed as described previously (Laha et al., 2015). BLAST search analyses (https://blast.ncbi.nlm.nih.gov/Blast.cgi?PAGE=Proteins) were performed using the full-length sequence of AtIPK2α retrieved from *A. thaliana* genome database (https://www.arabidopsis.org/).The retrieved sequences were filtered to include only those sequences with a percent identity of more than 30%, query cover of more than 35%, E-value of less than 10^-5^ and a bit score of more than 80. Sequences from every species were screened for the presence of gene isoforms. The GUIDANCE2 server (http://guidance.tau.ac.il/) was used to align the sequences using the version of MAFFT available on the server. The GUIDANCE2 algorithm was used to estimate the reliability of alignment columns. Unreliable columns having a confidence score lesser than 0.93 were removed from the multiple sequence alignment (GUIDANCE Server – a web server for assessing alignment confidence score (tau.ac.il)). Phylogenetic analysis was conducted in MEGA11(Tamura et al., 2021). The best maximum likelihood model for the given alignment was estimated using the default settings and this model was used to estimate a maximum likelihood tree. Alignment of complete protein sequences were done using ClustalW and phylogenetic tree was constructed with the Maximum Likelihood method, using the Dayhoff’s Model and a bootstrap test of 1000 replicates. Values less than 50% are not displayed on the tree and branch lengths are given in terms of the expected number of amino acid substitutions per site.

### Plant materials and growth conditions

Seeds of *Arabidopsis thaliana* wild-type (accession Col-0) and T-DNA insertional mutants of *ipk2β-1* (SALK_104995) and *ipk2α-1* (GABI_879D07) were genotyped for homozygous T-DNA insertions using primer Supplementary Data Set 1. Surface sterilization of seeds was performed by incubating the seeds in solution containing 0.05% SDS in 70% ethanol for 15 min and cold-stratified for 3 days and then germinated on solidified half-strength of Murashige and Skoog (½ MS) media. Growth chambers (Percival) were maintained at 22°C with long-day (LD) conditions (16 h:8 h; light: dark cycle) with light intensity 100 μmol/m^2^/s. The germinated seedlings were transferred onto soil (composition perlite and solite in the ratio of 1:2). The growth room condition was maintained at 22°C with 70% RH and long-day (LD) conditions (16 h:8 h; light: dark cycle) with light intensity 100 μmol/m^2^/s. *Marchantia polymorpha* accession Takaragaike-1 (Tak-1, male accession) was used as wild-type plants. *M. polymorpha* lines were propagated asexually by gemma cultured on half-strength Gamborg’s B5 medium with 1% phytagel under continuous light (50-60 µmol/m^2/^s) in a Percival growth chamber at 22⁰C.

### Molecular cloning

Cloning of *AtIPK2α* and Mp*IPMK in pET28b-His*_8_*-MBP* bacterial and yeast expression vector. Full-length coding sequence of AtIPK2α and MpIPMK were amplified using cDNA prepared from total RNA extracts of Col-0 and Tak-1 plants, respectively with primers listed in Supplementary Data Set 1. The amplified product was cloned at the EcoRV site in pBLUESCRIPT via blunt end cloning followed by directional subcloning into pET28a between the BamHI and Not1I sites. PCR mutagenesis was used to generate a point mutation in the conserved PXXXDXKXG motif of *AtIPK2α* and Mp*IPMK.* Primer used to generate side directed mutagens are enlisted in Supplementary Data Set 1. For cloning Mp*IPMK* wild-type and catalytic dead variants in yeast expression vector pCA45, the amplified product of Mp*IPMK* CDS from Tak-1 plants was subcloned with pBLUECRIPT followed by directional cloning in pCA45 vector pre-digested with the BamH1 and Not1.

### Protein expression and purification

To purify AtIPK2α protein, *E. coli* BL21 (RIL) strain was transformed with the pET28b-His_8_-MBP-AtIPK2α vector. A single colony was used to inoculate in terrific broth (TB medium). The protein induction was with 0.5 mM IPTG, the culture was incubated further for 3 days at 12⁰C. Cells were harvested at 6000 rpm for 10 min at 4⁰C and pellet was washed with lysis buffer (300 mM NaCl, 25 mM Na_2_HPO_4_, pH 7.5). The pellets were resuspended in lysis buffer having 5 mM β-ME and 1 mM PMSF, followed by cell lysis using sonication at 10 pulse of 30 sec ‘ON’ and 10 sec ‘OFF’. After sonication, the lysates were centrifuged at high speed 18000 rpm for 45 min at 4⁰C. Meanwhile Ni-NTA resin (Qiagen) was washed with ultra-pure water and equilibrated with PBS buffer twice. After centrifugation, the cleared protein supernatant was incubated with prewashed Ni-NTA beads for 6 h on rotor at 4⁰C. After incubation of the beads with protein supernatant, the beads were washed thrice with wash buffer (lysis buffer, 5 mM β-ME, 10 mM imidazole) at 4000 rpm for 5 min at 4⁰C. For elution, the beads were incubated with 250 µL elution buffer (lysis buffer containing 5 mM β-ME, 250 mM imidazole buffer) at 4⁰C for 5 min on a rotor and centrifuged at 4000rpm for 5mins. Three such elutions were collected. Aliquots of elution and different concentration of BSA standards were heated at 95⁰C after adding 1X SDS loading dye and loaded on 12% SDS-PAGE. Protein was visualized by coomassie staining and quantification of the band intensity was done using ImageJ. The mutant variants of AtIPK2α purification were performed as mentioned above. MpIPMK protein and its catalytic dead variants were purified using PBS buffer (137 mM NaCl, 2.7 mM KCl, 10 mM Na_2_HPO_4_, 1.8 mM KH_2_PO_4_).

### *In vitro* kinase assay

The InsP_6_ kinase assays were performed by incubating 5 µg of recombinant protein of AtIPK2α and AtIPK2β and their catalytic dead variants in a 30 µL of reaction mixture containing kinase buffer [5 mM MgCl_2_, 20 mM HEPES (pH 7.5), 1mM DTT], 12.5 mM ATP, and 10 nmol InsP_6_ at 37⁰C for 12 h. MBP protein was used as negative control and AtITPK1 (Laha et al., 2019) was used as positive control were added in equal concentration. The *in vitro* kinase assay of MpIPMK and its catalytic dead variants was performed using the above-mentioned reaction condition. The reaction products were resolved using PAGE(Pisani et al., 2014) and visualized by toluidine blue staining.

### Enrichment of inositol phosphate using titanium dioxide bead

Inositol phosphate pull down was performed as described previously (Laha et al., 2019; Wilson et al., 2015). All steps were carried on ice. TiO_2_ beads were weighed to 4-5 mg for each sample and washed twice in water and once in 1 M perchloric acid (PA). Liquid N_2_-frozen plant material (approx. ∼ 100 mg) was homogenized using a pestle and immediately resuspended in 800 µL ice-cold PA. Samples were kept on ice for 5 min with short intermediate vortexing and then centrifuged for 5 min at 20000 g at 4⁰C. The supernatants were transferred into fresh 1.5 mL tubes and centrifuged again for 5 min at 20000 g. The supernatants were resuspended in the prewashed TiO2 beads and incubated at 4⁰C for 30 min. After incubation, the beads were pelleted by centrifuging at 8000 g for 1 min and washed twice in PA. The supernatants were discarded. To elute inositol polyphosphates, beads were resuspended in 200 µL 10% ammonium hydroxide and then incubated for 5 min at room temperature. After centrifuging at 2600g, the supernatants were transferred into fresh tubes. The elution process was repeated, and the second supernatants were collected as well. Eluted samples were vacuum evaporated at room temperature to dry completely. InsPs were resuspended in 40 µL ultra-pure water for the following CE-MS analysis.

### CE-MS analysis

The analysis was performed on a CE-QQQ system (Agilent 7100 CE-with Agilent 6495C Triple Quadrupole and Agilent Jet Stream electrospray ionization source, adopting an Agilent CE-ESI-MS interface). An isocratic Agilent 1200 LC pump was used to deliver the sheath liquid (50% isopropanol in water) with a final splitting flow speed of 10 µl/min via a splitter. All separation was performed via a bare fused silica capillary with a length of 100 cm (50 µm internal diameter and 365 µm outer diameter). 40 mM ammonium acetate titrated with ammonium hydroxide to pH 9.0 was used as background electrolyte (BGE). Between runs of each sample, the capillary was flushed with BGE for 400s. Samples were injected by applying 100 mbar pressure for 15s (30 nL). The MS source parameters were as follows: gas temperature was 150°C, gas flow was 11 L/min, nebulizer pressure was 8 psi, sheath gas temperature was 175°C and with a flow at 8 L/min, the capillary voltage was –2000V, the nozzle voltage was 2000V. Negative high pressure RF and negative low pressure RF were 70 V and 40 V, respectively. Multiple reaction monitoring (MRM) transitions were setting as shown in Supplementary Data Set 6.

Internal standard (IS) stock solution of 8 µM [^13^C_6_] 2-OH InsP_5_, 40 µM [^13^C_6_] InsP_6_, 2 µM [^13^C_6_] 1-InsP_7_, 2 µM [^13^C_6_] 5-InsP_7_, 1 µM [^18^O_2_] 4-InsP_7_ (only for assignment of 4/6-InsP_7_) and 2 µM [^13^C_6_] 1,5-InsP_8_ were spiked to samples for the assignment of isomers and quantification of InsPs and PP-InsPs. 5 µL of the IS stock solution was mixed into 5 µL samples. Quantification of InsP_8_, 5-InsP_7_, 1-InsP_7_, InsP_6_, and InsP_5_ was performed with known amounts of corresponding heavy isotopic references spiked into the samples. Quantification of 4/6-InsP_7_ was performed with [^13^C_6_] 5-InsP_7_ and Quantification of InsP_3_ and InsP_4_ of which no isotopic standards are available was performed with spiked [^13^C_6_] InsP_6_. After spiking, 4 μM [^13^C_6_] 2-OH InsP_5_, 20 μM [^13^C_6_] InsP_6_, 1 μM [^13^C_6_] 5-InsP_7_, 1 μM [^13^C_6_] 1-InsP_7_, and 1 µM [^13^C_6_] 1,5-InsP_8_ were the final concentration inside samples. All [^13^C_6_] inositol references (Harmel et al., 2019) were kindly provided by Dorothea Fiedler.

### Yeast strains and transformation

The yeast strains were grown on YPD agar plates at 28⁰C. Transformed yeast strains were grown in complete minimal medium containing the appropriate amino acids, 2% glucose, and lacking uracil to maintain selection for URA3 plasmids at 28⁰C. Yeast transformations were performed by the lithium acetate method (Gietz et al., 1992). Briefly, single colony of streaked yeast strain was inoculated in 5 mL of YPD liquid medium and incubated at 28⁰C in shaker incubator for overnight. Fresh 4 mL YPD liquid media was inoculated with overnight grown culture to reach OD_600nm_ ∼0.6. And culture was allowed to grow for 4 h in shaker incubator. Once the culture reached OD∼ 1, the culture was harvested at 2600 rpm for 1 min at room temperature. The pellets were washed twice with 500 µL of TE/LiAc buffer at 2600 rpm for 2 min. After final wash, the cells were resuspended with 200 µL of TE/LiAc buffer and kept on ice. These yeast competent cells were used for transformation. A total of 3.5 µL of salmon sperm DNA (approx. 8 mg/mL) was heated at 95⁰C for 5 min and kept on ice for 2 min. 1 µL of plasmid (200 – 500 ng plasmid) was added to the salmon sperm DNA followed by adding 16.5 µL of yeast competent cells. The cells with plasmid were resuspended with PEG/LiAc buffer and incubated for 40 min at room temperature on rotor. After incubation, the cells were subjected to heat shock at 42⁰C for 20 min. 70 uL of the cell suspension was used for plating on selection plate. The plates are incubated at 28⁰C. Complementation assay was performed by dropping 100-fold serially diluted resuspension of transformed colony of yeast on selection and screening plate.

Supplementary Data Set 4. contains the list of yeast strains used in the experiment of complementation assay.

### RNA extraction, cDNA synthesis and gene expression analyses

RNA extraction and cDNA synthesis was done as described previously(Pullagurla et al., 2023). Briefly, 100 mg of plant tissue was used for RNA extraction with TRIzol (Sigma Aldrich) reagent, followed by DNase treatment with DNase I (NEB, M0303). A total of 2-3 µg of RNA was used for cDNA synthesis using PhiScript™ cDNA Synthesis Kit (dx/dt). The qPCR was performed using the DyNAmo ColorFlash SYBR Green qPCR Kit (Thermo-scientific) with CFX96 Touch Real-Time PCR Detection System (Bio-Rad Hercules, CA, USA) according to the manufacturer’s protocol (Bio-Rad). Relative expression was calculated according to relative quantitation method (ΔΔCT). *ACTIN2* and Mp*ACT7* were used as reference genes for the qPCR analysis in *A. thaliana* and *M. polymorpha*, respectively. The primers used for qPCR analyses are detailed in Supplementary Data Set 1.

### Extraction and HPLC analysis of inositol phosphates

Inositol polyphosphates were extracted from yeast and analyzed as described (Azevedo and Saiardi, 2006; Laha et al., 2015; Laha et al., 2021a). Yeast transformants were grown to midlog phase in minimal media, labelled in 2 mL of minimal media supplemented with 6 μCi/mL [^3^H]-*myo*-inositol. The cells were harvested, washed twice with ultra-pure water were extracted in 1 M perchloric acid extracted. Extracted inositol phosphates from yeast were resolved by strong anion exchange high performance liquid chromatography (SAX-HPLC) using a Partisphere SAX 4.6 x 125 mm column (Whatman) at a flow rate of 0.5 mL/min with a shallow gradient formed by buffers A (1 mM EDTA) and B [1 mM EDTA and 1.3 M (NH_4_)_2_HPO_4_, pH 3.8, with H_3_PO_4_] (Azevedo and Saiardi, 2006).

### Cloning and plasmid construction for plant transformation

Mp*IPMK* guideRNA (gRNA) designing and cloning. CRISPR/Cas9-based genome editing of Mp*IPMK* was performed as described previously(Sugano et al., 2018; Sugano et al., 2014). Selection of the guideRNA (gRNA) target site was done using CRISPRdirect web tool (Naito et al., 2015). The gRNA protospacers were generated by annealing complementary oligonucleotides and inserted into the pMpGE_En03 vector(Sugano et al., 2018) previously digested with BsaI. Mp*IPMK*-gRNA was incorporated into the binary vector pMpGE010 (Sugano et al., 2018) using Gateway LR Clonase II Enzyme mix (Invitrogen, 11791100). In total, eight gRNAs were designed at different part of the gene that were used for generating CRISPR lines.

### Thallus transformation of *M. polymorpha*

Transformation of *M. polymorpha* was performed as described previously(Sugano et al., 2014). In short, 14-day-old gemmalings were sliced to eliminate the apical notches and kept for 3 days for regeneration on half strength Gamborg’s B5 medium supplemented with 1% sucrose. Regenerating thalli fragments were then co-incubated with *Agrobacterium tumefaciens GV3010* cells carrying the corresponding vectors in half strength Gamborg’s B5 medium supplemented with 2% sucrose under white light and gentle shaking of 130 rpm at 22⁰C. After three days of co-culture, the plant fragments were washed three times with sterile water and placed on half strength Gamborg’s B5 medium supplemented with 100 µg/mL cefotaximine and 10 µg/mL hygromycin B.

### Construction of RNAi plasmid and generation of knockdown lines of in *A. thaliana*

Full-length coding region of AtIPK2α (∼900bp) was cloned into pOpOff2(Hyg)(Parvin et al., 2017) by Gateway recombination (LR ClonaseII, Invitrogen). The resulting vector pOpOFF2(Hyg)-AtIPK2α was used to stably transform the *atipk2β* knockout plants using *Agrobacterium tumefaciens* GV3101-mediated transformation by floral dipping method as described previously (Laha et al., 2015). Positive transformants were selected on ½ MS agar plates containing 30 mg/mL hygromycin. Homozygous lines were identified from selected lines at the T3 generation. For induction of RNAi, plants were treated with 25 µM dexamethasone (Dex) as indicated for each experiment.

### Thermotolerance assay

Thermotolerance assays were performed as described previously (Guan et al., 2013; Ibanez et al., 2018; Li et al., 2023; Zhang et al., 2021). For basal thermotolerance assay, 7-day-old seedlings of all the genotype (Col-0, *atipk2α*, *atipk2β* and *atipk2β^-/-^atipk2α^kd^*) grown on ½ MS medium were exposed to 37⁰C for 2½ days and were kept at 22⁰C for recovery of another 4 days. Photographs were recorded and survival rates were counted after 4 days of recovery. For root thermomorphogenesis study, the stratified seedlings were grown at 22⁰C on ½ MS medium for 4 days, after which they were transferred to 28⁰C or maintained at 22⁰C for 5 days. Photographs were taken after 5 days of transfer and root length measurement was performed using ImageJ software. For hypocotyl elongation assay, the stratified seedlings were allowed to grow at 22⁰C on ½ MS medium and then transferred to 28⁰C or maintained at 22⁰C for another 4 days. Photographs were taken after 4 days of transfer and the hypocotyl length was measured using Image software. For thermotolerance assay in *M polymorpha*, gemma of WT and Mp*ipmk* mutant plants were transferred on half strength Gamborg’s B5 containing 1% agar plates and were allowed to grow at 22⁰C with 16 h:8 h; light: dark cycle for 5 days. On the 5^th^ day, the plates were placed in water bath pre-heated at 37⁰C for 8 h. The heat-treated plates were then placed back at 22⁰C chamber for recovery. Images were recorded after 4 days of recovery. Survival of the thallus were assessed by their ability to grow and maintain green thallus.

For thermomorphogesis study in *M. polymorpha*, gemmae of WT and Mp*ipmk* knockout lines were transferred on half strength Gamborg’s B5 media with 1% agar and grown for 5 days at 22⁰C with white light of (16 h:8 h; light: dark cycle) for 5 days. On the 5^th^ day, the plates having WT and Mp*ipmk* knockout lines were exposed to 37⁰C in a plant chamber. Plates were exposed to heat shock for different time points, 6 h, 8 h and 10 h. The heat-exposed plants were kept back at 22⁰C. Images were taken after 14 days of recovery. Thallus area was calculated using ImageJ software. Data were analysed using GraphPad Prism 6.

### GUS staining

The 14-day-old seedlings of Col-0 and *atipk2β^-/-^atipk2α^kd^* lines were fixed for 30 min in ice-cold 90% (v/v) acetone and rinsed with staining buffer (0.5 M sodium phosphate buffer pH 7.2, 10% Triton X, 10 100 mM potassium ferrocyanide, 100 mM potassium ferricyanide) without X-Gluc (500 µg ml−1 5-bromo-4-chloro-3-indolyl-β-D-glucuronide) followed by incubation in staining buffer with 2 mM X-Gluc at 37 °C in the dark for overnight. After a series of wash with 20%, 35% and 50% ethanol, the cleared seedlings were mounted on slide and imaged using Zeiss microscope.

### Luciferase assay

A luciferase reporter system was used to study the transactivation activity. The effectors *AtIPK2α* and *HSFA1V* were cloned under constitutive promoter with N-terminal fused YFP and GFP tag, respectively. The promoter of *AtHSP18.2* was cloned in pGWB535 with gateway cloning method. Both effectors and reporter were transformed in *Agrobacterium* (GV3101) and infiltrated in *Nicotiana benthamiana* plants as described previously (Laha et al., 2015). Leaves were collected 72 h of infiltration. Luciferase assay was performed by luciferase assay kit (Promega; E152A) as manufacturer’s instruction. Bioluminescence was measured using GloMax Explorer multimode microplate reader (Promega).

### Statistical analysis

Statistical analyses were performed using GraphPad Prism 6 software. The details of statistical data have been provided in Supplementary Data Set 7.

### Accession numbers

*AtIPK2α* (At5g07370), *AtIPK2β* (At5g61760), At*ITPK1* (At5g16760), *Actin2* (At3g18780), *AtHSP22* (At4g10250), *AtHSP70* (At3g12580), *At*HSP90 (At4g24190), *AtHSP17.6* (At5g12020), Mp*IPMK* (Mp1g22660.1) and Mp*ACT7* (Mp6g11010)

## Acknowledgements

We thank Y. Hirakawa for providing us the *Marchantia polymorpha* strains. We acknowledge Cristina Azevedo for the pCA45 plasmid. We thank Saikat Bhattacharjee for providing us the Arabidopsis *ipk2α-1* seeds.

## Author contributions

R.Y. generated the transgenic lines, performed the phenotyping and biochemical experiments and gene expression analyses. G.L. performed the CE-MS analyses of the *Arabidopsis* and *M. polymorpha* transgenic lines. P.R. performed biochemical experiments. N.J.P. generated structural models and performed phylogenetic analysis. D.Q. performed CE-MS analyses of the *in vitro* reactions and plant samples. H.J.J. supervised the project. D.L. conceived, supervised, and designed the study. D.L. obtained the funding. R.Y., G.L., P.R., N.J.P., D.Q., H.J.J. and D.L. analyzed the data. R.Y. and D.L. wrote the manuscript with input and approval from all the authors.

