## Supplemental file for "Conservation of heat stress acclimation by the inositol polyphosphate multikinase, IPMK responsible for 4/6-InsP_7_ production in land plants"

### Supplementary Figures

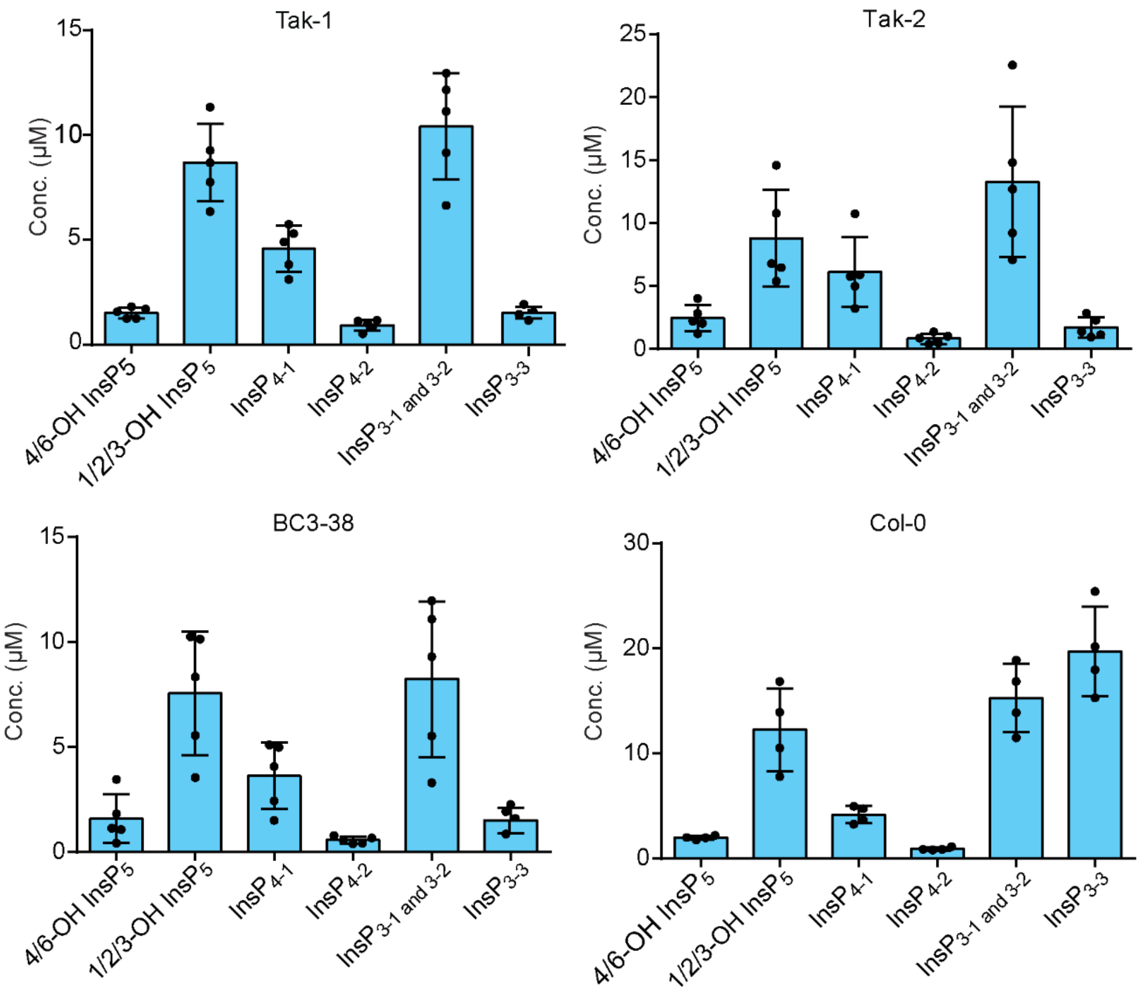


**Supplementary Figure 1. Profile of lower inositol phosphates in the Tak-1, Tak-2, BC3-38 *M. polymorpha* plants and Col-0 plants.**

Quantification of CE-MS analyses of different lower inositol phosphates and their isomers in the above-mentioned embryophytes. 14-day-old *M. polymorpha* thalli (Tak-1, Tak-2 and BC3-38) and 7-day-old *A. thaliana* (Col-0) seedlings were harvested, and inositol phosphates were extracted using TiO2-based pull down method (see details in the method section) and subjected to CE-MS. The $InsP_5$ species were assigned by mass spectrometry and identical migration time compared with relative standards. Two $InsP_4$ and three $InsP_3$ isomers were detected. Data are means ± SE (n = 5 biological replicates).

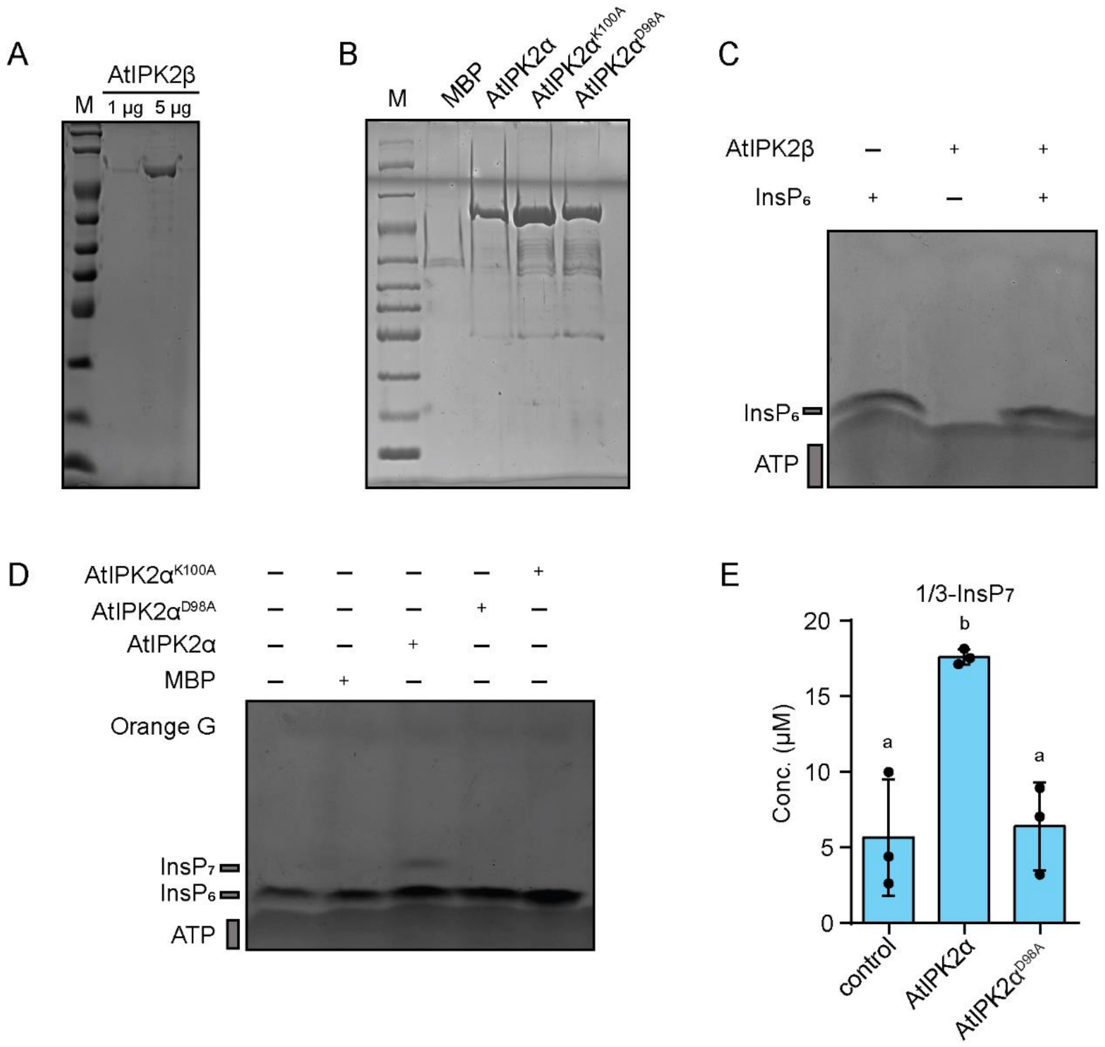


**Supplementary Figure 2. AtIPK2α phosphorylates $InsP_6$ *in vitro.***

**A** and **B**. SDS-PAGE analysis of AtIPK2β (A), WT and the catalytic dead variants of AtIPK2α (B). Resolved proteins were visualized by coomassie blue staining.

**C.** PAGE analysis of *in vitro* kinase assay reaction of AtIPK2β. Recombinant $His_8$-MBP-AtIPK2β was incubated with $InsP_6$ and ATP at 37ºC for 12 h. $InsP_6$ alone and AtIPK2β served as control. No reaction product could be detected using PAGE.

**D.** PAGE analysis of the *in vitro* kinase reaction products of AtIPK2α. Recombinant $His_8$-MBP-AtIPK2α and the catalytic dead proteins, AtIPK2α $^{D98A}$, AtIPK2α $^{K100A}$ were incubated with 12.5 mM ATP, and 10 nmol $InsP_6$ at 37ºC for 12 h in reaction buffer. The reaction product was separated by 33 % PAGE and visualized with toluidine blue. $His_8$-MBP served as a negative control.

**E.** Quantification of the AtIPK2α reaction product using CE-MS analyses. A minor amount of 1/3-Ins$P_7$ species could be detected in the reaction products. Data represent means ± SEM (n = 3). Letters depict the significance in one-way ANOVA followed by Dunnett's test ($P$ <0.05).

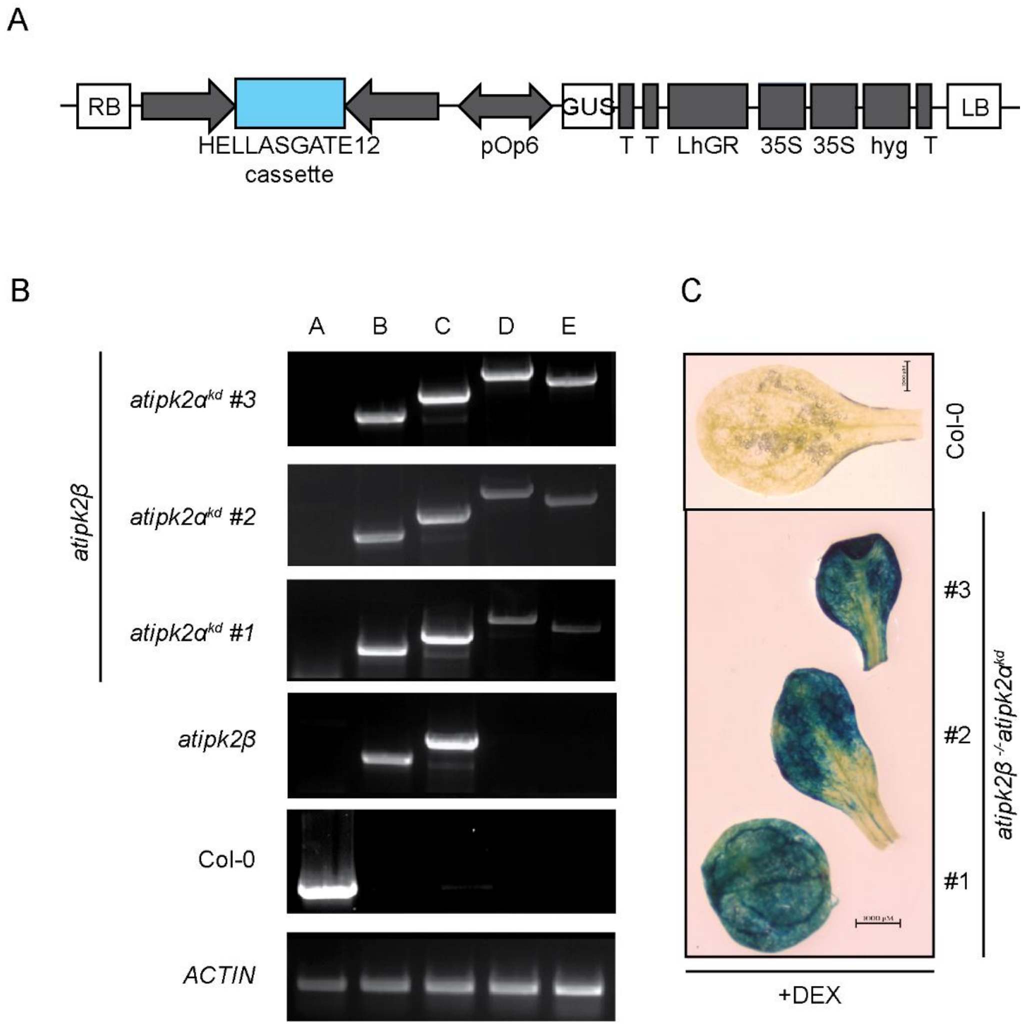


**Supplementary Figure 3. Generation of *atipk2α* knockdown lines in the *atipk2β* knockout plants.**

**A.** Schematic diagram of the pOpOff2 vector. RB, right border; T, terminator; hyg, hygromycin; LB, left border.

**B.** Genotyping PCR of Col-0, *atipk2β* and all the three *atipk2α* knockdown lines. A to E represents the primers set used for genotyping, details of the primers are mentioned Supplementary Data Set 1. *ACTIN* served as a reference gene.

**C.** Representative images of the GUS signal in leaves of Col-0, *atipk2β* and the three independent *atipk2β*$^{-/-}$ *atipk2α*$^{kd}$ lines used in this study after dexamethasone treatment.

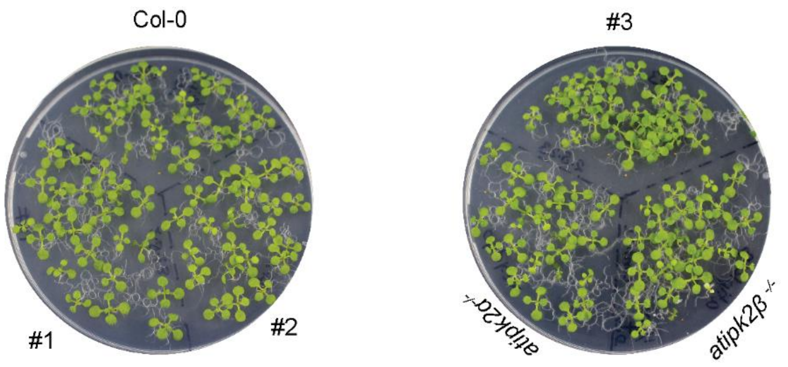


**Supplementary Figure 4. Profile of different inositol phosphates after heat shock.**
Photograph of the control plate maintained at 22ºC throughout basal thermal tolerance assay. This is the control set for the experiment presented in the main Figure 3C.

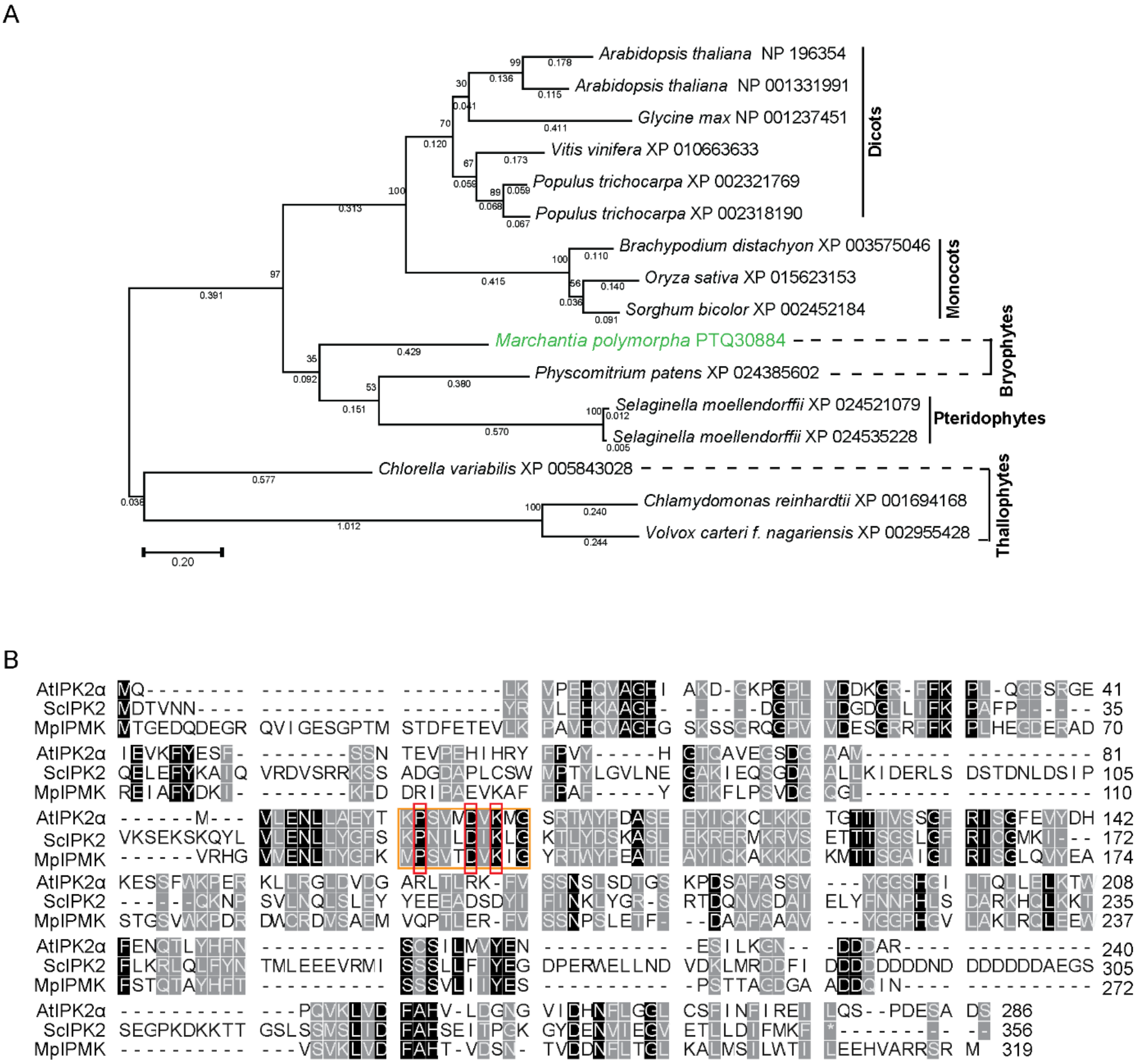


**Supplementary Figure 5. AtIPK2α-type kinases are ubiquitous in green plants and share conserved catalytic motif.**

**A.** The phylogenetic tree was estimated from an alignment of AtIPK2α amino acid sequences using maximum likelihood. Branch support was calculated from 1000 bootstrap replicates, and values below 50% are omitted. Branch lengths are given in terms of expected numbers of amino acid substitutions per site.

**B.** Protein alignment of MpIPMK with AtIPK2α and ScIPK2. Red rectangle marks the conserved catalytic motif PXXXDXKXG of the InsP kinase.

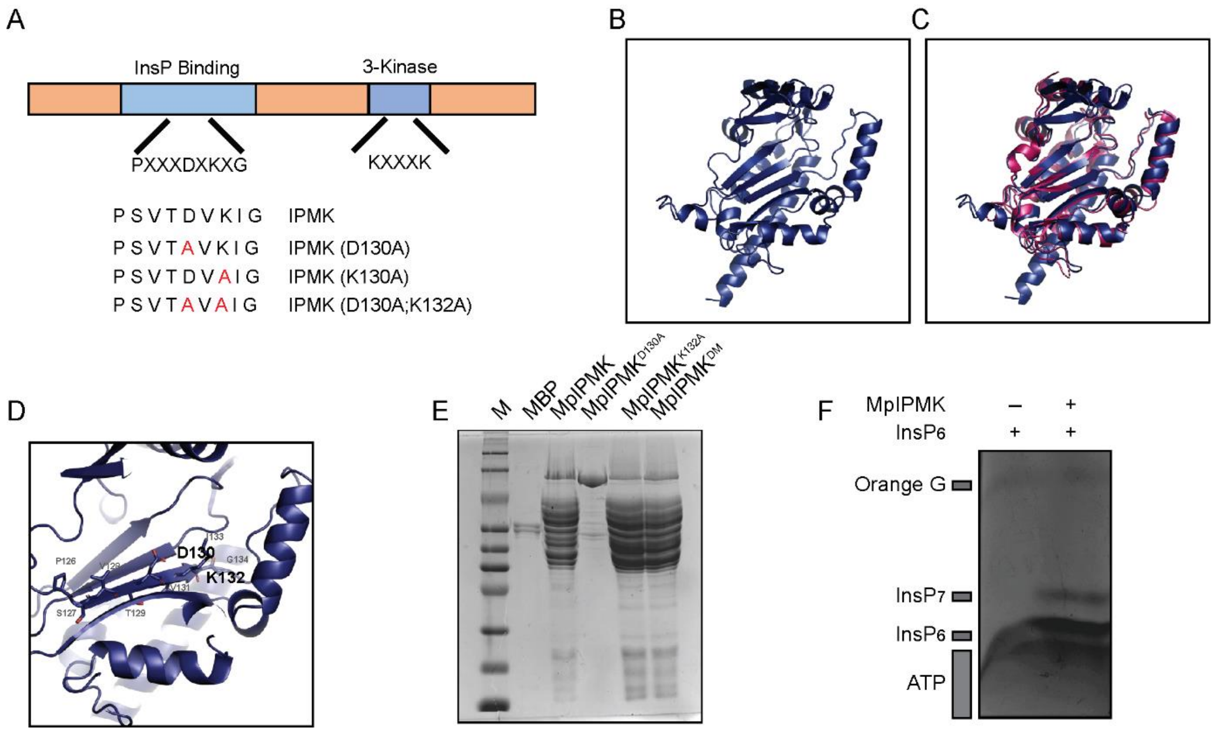


**Supplementary Figure 6. Structural model and activity of MpIPMK.**

**A**. Cartoon depicting the conserved PXXXDXVXG motif of IPMK-type proteins. The residues highlighted in red are the altered residues, forming catalytic dead variants of IPMK i.e., $MpIPMK^{D130A}$, $MpIPMK^{K132A}$, $MpIPMK^{D130AK132A}$ (referred as $MpIPMK^{DM}$).

**B.** Structural model (overview) of MpIPMK based on AtIPK2α (Protein Data Bank entry 4FRF). Models were obtained by the AlphaFold web portal (https://alphafold.ebi.ac.uk/) and built on the Pymol.

**C.** Structural overlay of AtIPK2α (hot pink), MpIPMK (blue) structures (RMSD value = 0.872). Note the similarity between the MpIPMK model and the AtIPK2α structure (Protein Data Bank entry 4FRF).

**D.** Zoom-in-into view of the catalytic active site of MpIPMK.

**E.** SDS-PAGE analysis of MpIPMK WT and catalytic dead variants.

**F.** PAGE analysis of the *in vitro* kinase reaction products of MpIPMK. Recombinant $His_8$-MBP-MpIPMK was incubated with 12.5 mM ATP, and 10 nmol $InsP_6$ at 37ºC for 12 h in reaction buffer. The reaction product was separated by 33 % PAGE and visualized with toluidine blue. $InsP_6$ alone served as a control.

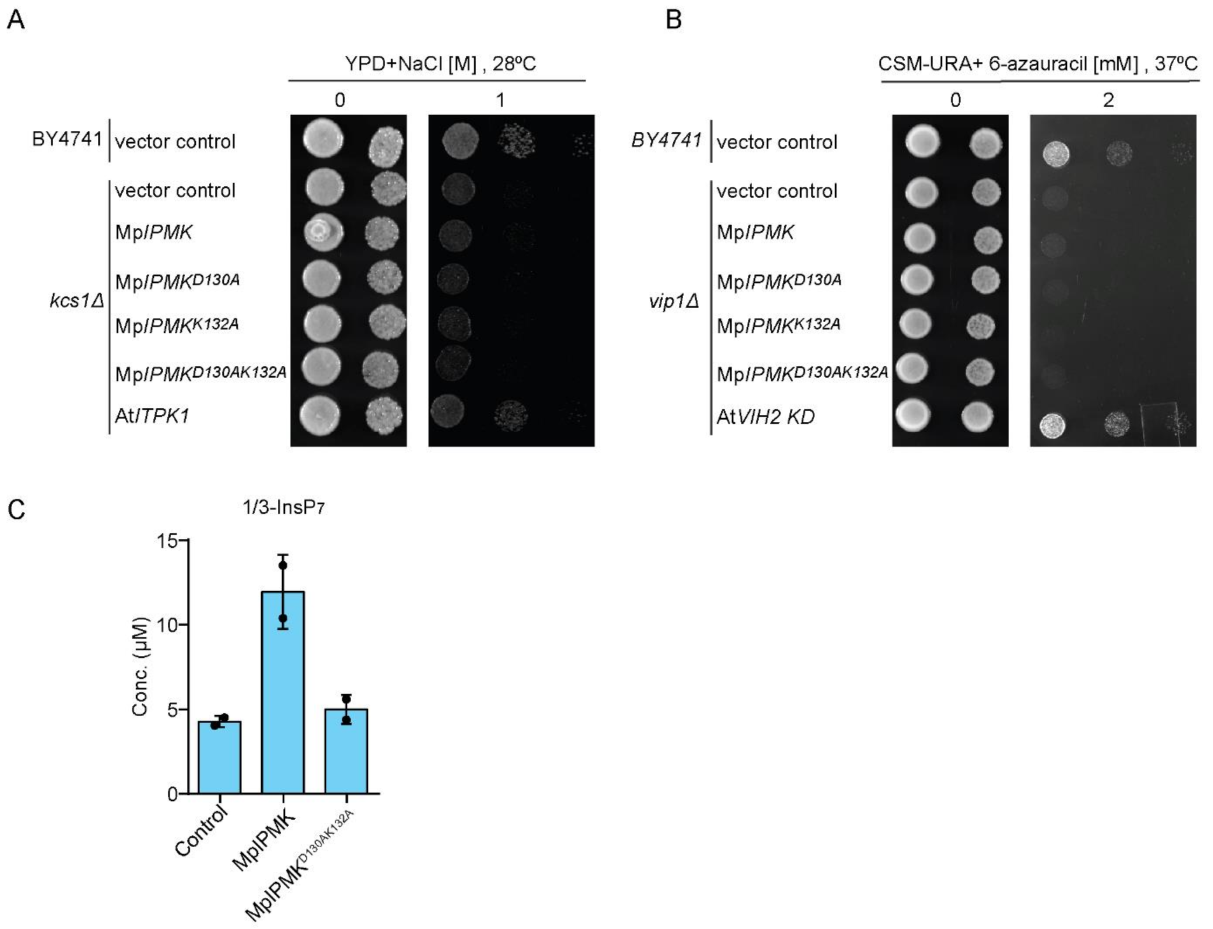


**Supplementary Figure 7. MpIPMK does not possess yeast Kcs1- or Vip1-like activity.**

**A.** Complementation of the yeast *kcs1*Δ-associated growth defects by the ectopic expression of Mp*IPMK*. Wild-type and *kcs1*Δ yeast transformants (BY4741 background) carrying designated plasmids were spotted in 8-fold serial dilution onto YPD with and without NaCl incubated at 28ºC and 37ºC. *AtITPK1* served as positive control[1] and empty vector served as negative control.

**B.** Complementation of *vip1*Δ -associated growth defects in yeast by ectopic expression of Mp*IPMK*. The *vip1*Δ yeast strain transformed with the episomal pCA45 (URA3) plasmids carrying Mp*IPMK* and kinase dead mutants were spotted in 8-fold serial dilutions onto uracil-free minimal medium in presence and absence of 6-azauracil. No rescue of phenotype was observed. *AtVIHKD* served as positive control[2] and empty vector served as negative control.

**C.** Quantification of the reaction product of MpIPMK analyzed by CE-MS. Data represent means ± SEM (n = 2).

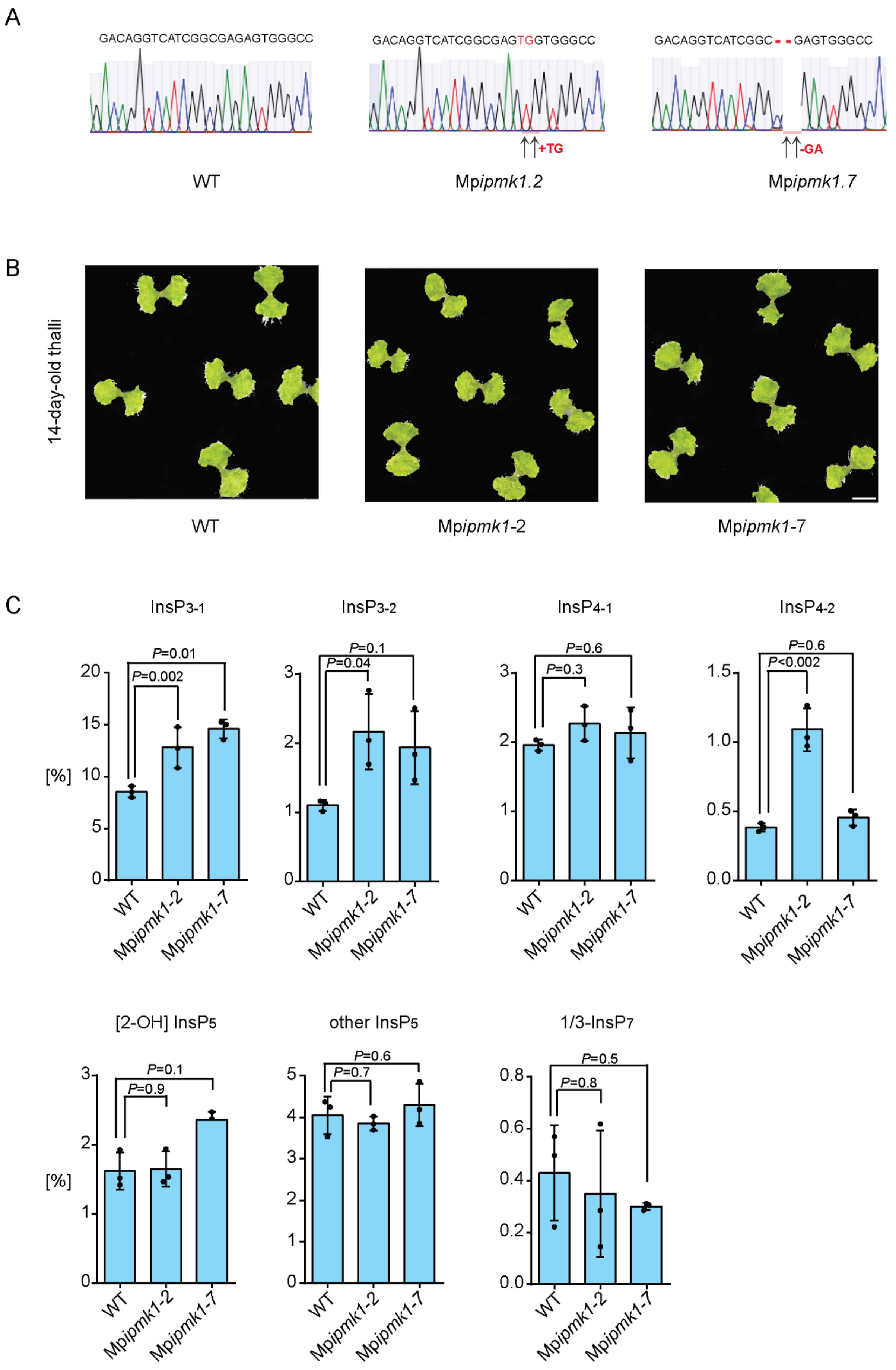
A
GACAGGTCATCGGCGAGAGTGGGCC
WT
GACAGGTCATCGGCGAGTGGTGGGCC
+TG
Mpipmk1.2
GACAGGTCATCGGC--GAGTGGGCC
-GA
Mpipmk1.7
B
14-day-old thalli
WT
Mpipmk1-2
Mpipmk1-7
C
InsP3-1
InsP3-2
InsP4-1
InsP4-2
[%]
P=0.01
P=0.002
P=0.1
P=0.04
P=0.6
P=0.3
P=0.6
P<0.002
WT
Mpipmk1-2
Mpipmk1-7
[2-OH] InsP5
other InsP5
1/3-InsP7
P=0.1
P=0.9
P=0.6
P=0.7
P=0.5
P=0.8

**Supplementary Figure 8. Mp*ipmk* knockout plants does not exhibit any obvious developmental defects.**

**A.** Chromatogram showing edited nucleotide sequences of Mp*ipmk1.2* and Mp*ipmk1.7* compared with those of WT plants using chromatogram obtained from sequencing results. **B.** Photograph of 14-day-old thalli of WT, Mp*ipmk1.2* and Mp*ipmk1.7* plants. Gemma of all the genotype were grown on half-strength Gamborg's B5 medium adjusted at pH 5.7 and supplemented with 1% Phyto Agar (Himedia), under continuous light (50-60 µmol/$m^{2/}s^2$) and long-day conditions (16 h light at 22ºC, 8 h dark at 20ºC) in plant chamber (Percival).

**C.** CE-MS analyses of different inositol phosphates isomers in WT and Mp*ipmk* knockout plants. The $InsP_5$ and $InsP_7$ species were assigned by mass spectrometry and identical migration time compared with their relative standards. Two $InsP_4$ and two $InsP_3$ isomers were detected. Data are means ± SE (n = 3, biological replicates). Significant difference is determined by one-way ANOVA followed by Dunnett's test.
